# Graph construction method impacts variation representation and analyses in a bovine super-pangenome

**DOI:** 10.1101/2022.09.17.508368

**Authors:** Alexander S. Leonard, Danang Crysnanto, Xena M. Mapel, Meenu Bhati, Hubert Pausch

## Abstract

Several models and algorithms have been proposed to build pangenomes from multiple input assemblies, but their impact on variant representation, and consequently downstream analyses, is largely unknown. We create multi-species “super-pangenomes” using pggb, cactus, and minigraph with the *Bos taurus taurus* reference sequence and eleven haplotype-resolved assemblies from taurine and indicine cattle, bison, yak, and gaur. We recover 221k nonredundant structural variations (SVs) from the pangenomes, of which 135k (61%) are common to all three. SVs derived from assembly-based calling show high agreement with the consensus calls from the pangenomes (96%), but validate only a small proportion of variations private to each graph. Pggb and cactus, which also incorporate base-level variation, have approximately 95% exact matches with assembly-derived small variant calls, which significantly improves the edit rate when realigning assemblies compared to minigraph. We use the three pangenomes to investigate 9,566 variable number tandem repeats (VNTRs), finding 63% have identical predicted repeat counts in the three graphs, while minigraph can over or underestimate the count given its approximate coordinate system. We examine a highly variable VNTR locus and show that repeat unit copy number impacts expression of proximal genes and non-coding RNA. Our findings indicate good consensus between the three pangenome methods but also show their individual strengths and weaknesses that need to be considered when analysing different types of variants from multiple input assemblies.

## Background

Pangenomes store and represent sequences from multiple individuals and enable unbiased variation-aware sequence variant analyses [1]. Graph pangenomes represent alleles that differ between the input assemblies as nodes (representing sequence) connected by edges [2]. Several methods have been proposed to construct graph pangenomes from genome-scale data. For instance, minigraph applies approximate mapping to construct structural variant-based pangenomes from multiple input assemblies [3]. The reference backbone has an impact on these pangenomes because it propagates bias when large segments are missing in the backbone assembly [4]. Recently, cactus [5] and pggb (pangenome graph builder) [6] have been proposed to construct pangenomes from multiple input assemblies using reference-free base-level alignment. The pggb and cactus pangenomes contain all types of differences found between the assemblies, ranging from single nucleotide to large structural differences with nested variation [2].

Advancements in long-read sequencing and algorithms enable automated assembly of reference-quality genomes also for species with gigabase-sized genomes and so request for pangenomes that seamlessly accommodate and represent an increasing number of genomic resources [7],[8]. For instance, the Telomere-to-Telomere (T2T) consortium recently reported on a first complete human genome assembly [9] but routine near-T2T assembly [10] is becoming increasingly possible in humans [11] and other vertebrate species [12]. The Human Pangenome Reference Consortium (HPRC) coordinates global sequencing and assembly efforts with the goal to build a community-accepted variation-aware pangenome that integrates multiple input assemblies and represents global human genomic diversity [8]. An initial analysis of pangenomes constructed from 47 phased diploid genome assemblies has recently been reported by the HPRC [13].

Likewise, a Bovine Pangenome Consortium (BPC) has been formed to coordinate assembly efforts for the global cattle genomics community (https://bovinepangenome.github.io/). High nucleotide diversity and the separation of millions of global cattle into several hundred distinct breeds with unique genetic features, as well as frequent hybridization with their undomesticated relatives, make cattle an appealing species to improve assembly techniques [14] and investigate pangenome construction [4]. Bovine pangenomes constructed from multiple reference-quality assemblies revealed that the linear *Bos taurus taurus* reference sequence lacks millions of bases that are accessible in assemblies from other individuals [4],[15],[7]. However, the impacts of different construction methods on pangenome profiles, variant representation, and downstream analyses are more uncertain particularly when they include assemblies from multiple species.

Here, we apply cactus, minigraph, and pggb to build multi-species super-pangenomes with the *Bos taurus taurus* reference sequence and eleven haplotype-resolved assemblies from taurine and indicine cattle, bison, yak, and gaur. We assess the properties of the resulting pangenomes, recover large and small variants, and investigate how the pangenomes represent different types of DNA variation. We then profile variable number tandem repeats (VNTR) to investigate how pangenomes integrate DNA variants that are challenging to resolve from linear alignments. Finally, we exploit phylogenetic relationships between the input assemblies to identify a highly polymorphic VNTR locus which mediates the expression of proximal genes and non-coding RNA.

## Results

We built pangenomes from autosomal sequences of domestic cattle and three of their wild relatives using minigraph, cactus, and pggb to assess how different graph pangenomes represent the same underlying input sequences. Nineteen haplotype-resolved assemblies from eight breeds of taurine (*Bos taurus taurus*) and indicine (*Bos taurus indicus*) cattle, yak (*Bos grunniens*), bison (*Bison bison bison*), and gaur (*Bos gaurus*) as well as the current Hereford-based *Bos taurus taurus* reference genome sequence were considered (**Figure 1a**), of which 12 were used for pangenome construction and 8 held out for analysis. These high quality genomes were assembled using different sequencing and algorithmic approaches, but this has limited effect on the pangenomes [7]. However, the larger amount of centromeric/telomeric sequence in the HiFi-based assemblies (**Figure 1b,c**) does require additional consideration. Minigraph used ARS-UCD1.2 as a backbone, whereas pggb and cactus are reference-free methods, although cactus is guided by an approximate phylogenetic tree (black subset of **Figure 1d**).

**Figure 1:**
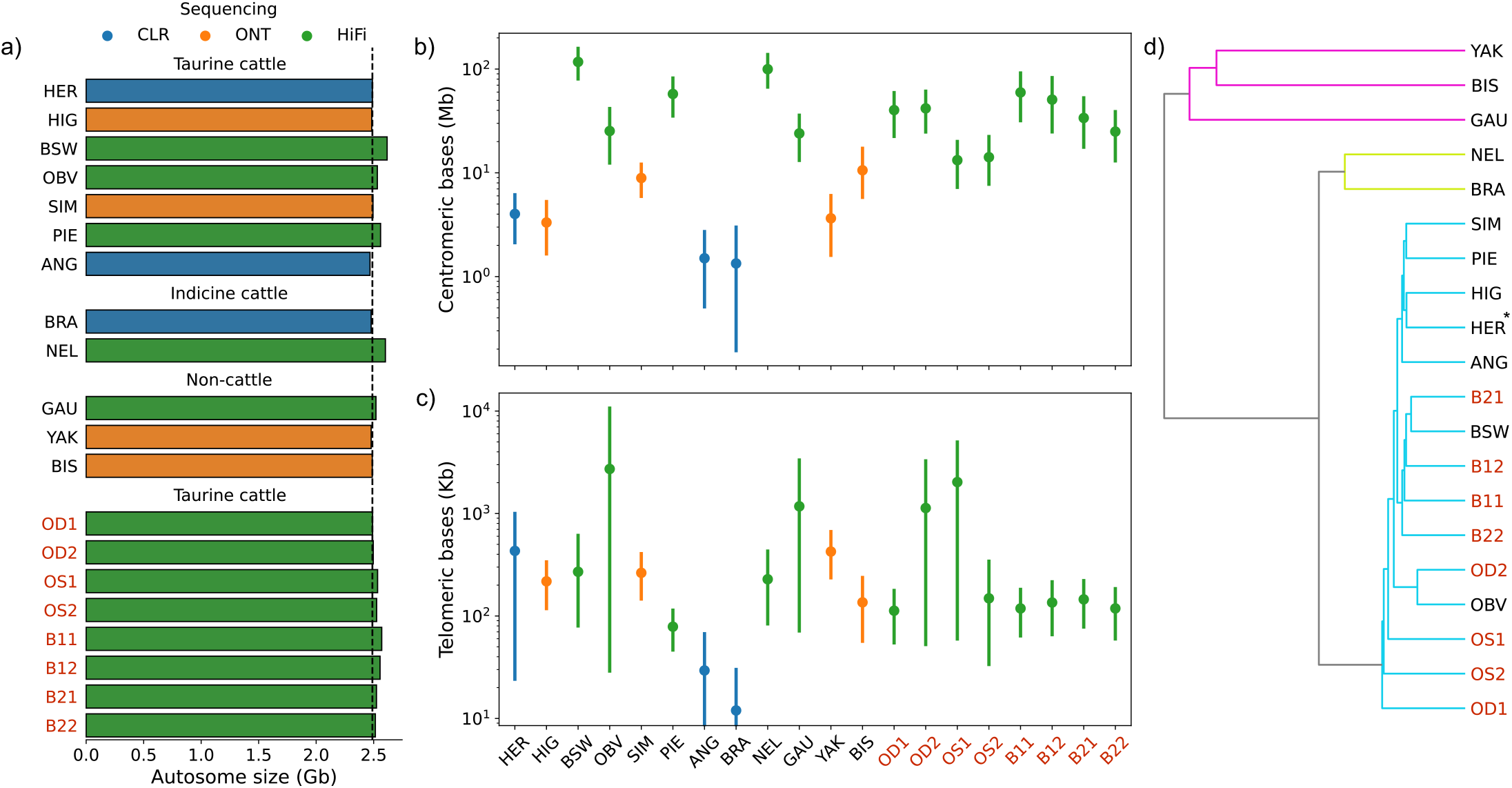
Input assemblies considered for the pangenome analyses. a) Autosomal length of the *Bos taurus taurus* reference sequence (HER, ARS-UCD1.2) and 19 haplotype resolved assemblies considered during pangenome construction and downstream analyses. Three pangenomes were created with ARS-UCD1.2 [HER] and eleven input assemblies (Brown Swiss [BSW], Piedmontese [PIE], Highland [HIG], Angus [ANG], Original Braunvieh [OBV], Simmental [SIM], Brahman [BRA], Nellore [NEL], gaur [GAU], bison [BIS], yak [YAK]), whereas eight additional Original Braunvieh or Brown Swiss assemblies indicated with red text (OD1, OD2, OS1, OS2, B11, B12, B21, B22) were only considered for downstream analyses. The colour of the bars indicates the primary sequencing technology used to construct the assemblies. The black dashed line indicates the length of ARS-UCD1.2. b) Centromeric and c) telomeric completeness is generally higher in the HiFi- and ONT-than CLR-based input assemblies. The marker is the sum over the autosomes and error bars indicate the 95% confidence interval. d) A tree constructed with mash from the assemblies reveals the expected separation between taurine, indicine and non-cattle. The * indicates the backbone genome used by minigraph.

### Pangenome construction and sequence content

The three pangenomes spanned between 427k and 198M nodes and contained between 2.6 and 3.0 gigabases (**Table 1**). Minigraph primarily incorporates larger (>50 bp) DNA variation, and so builds a smaller graph compared to pggb and cactus which include all sizes of variation, which is also reflected in the number of path steps in the graph and final file sizes. The non-reference nodes contain nearly five times as much sequence in cactus (552 Mb) and pggb (523 Mb) than minigraph (109 Mb). Pangenome construction took 16- and 253-times more CPU hours and required 8- and 7-times more memory for cactus and pggb than minigraph.

**Table 1.** Profiles of three bovine pangenomes constructed from autosomal sequences of twelve assemblies. Path steps are the total number of steps needed to trace all 12 assemblies through the graph. * Minigraph required an additional 18 CPU hours to add P-lines. † Cactus uses soft masked assemblies as input, which required an additional 1571 CPU hours.

| Parameter | Unit | minigraph | cactus | pggb |
| --- | --- | --- | --- | --- |
| Nodes | N | 427,012 | 198,431,246 | 179,575,371 |
| Edges | N | 606,926 | 272,102,708 | 245,150,846 |
| Node length | bp | 2,598,811,581 | 3,041,026,095 | 3,012,039,323 |
| Path steps | N | 3,358,976 | 1,621,936,527 | 1,442,793,659 |
| Repetitive sequence | bp | 1,107,501,421 | 1,361,489,638 | 1,415,552,890 |
| Centromeric sequence | bp | 2,939,789 | 291,982,193 | 255,091,362 |
| CPU time | h | 14* | 226† | 3,559 |
| Max memory | GiB | 7 | 54 | 46 |
| GFA file size | GB | 2.6 | 26.1 | 23.7 |

Pggb and cactus also contain more repetitive sequence than minigraph, 46.4%, 44.3%, and 42.2% respectively, although this is largely due to including centromeric sequence. The ARS-UCD1.2 reference genome contains little centromeric sequence, preventing minigraph from anchoring centromeric sequence present in the HiFi-based (Brown Swiss, Nellore, Original Braunvieh, Piedmontese, and gaur) and to a lesser extent in ONT-based (bison and Simmental) assemblies into its pangenomes. Although pggb and cactus do include centromeric sequence into their pangenomes, they span a similar amount of bases as the sum of the input assemblies, suggesting either they are biologically distinct or cannot be effectively collapsed into a graph representation in a way similar to other repeat elements (e.g. SINE, LINE, LTR, etc., **Supplementary Figure 1**).

### Consensus of genomic variation

To assess the commonality of genomic variation across pggb, cactus, and minigraph, as well as the suitability of their representation for downstream analyses, we decomposed and normalized the graph pangenomes into VCF files with respect to the ARS-UCD1.2 reference. Decomposing default minigraph pangenomes failed to output many expected structural variant (SV) alleles, particularly deletions in multi-node bubbles (**Supplementary Figure 2**). By adding path information (P-lines in GFA) through minigraph-based realignment, we recovered an additional 21,770 SVs (13.5% increase). Realignment occasionally produces paths incongruent with graph topology, where an assembly does not trace through nodes originating from that same assembly, but only affects approximately 0.03%, 0.03%, and 0.05% of taurine, indicine, and non-cattle nodes (**Supplementary Figure 3**). Cactus had the most SV genotypes marked as CONFLICT (2.2%), indicating that a haploid assembly had multiple possible ambiguous alleles, with minigraph and pggb significantly lower, 0.5% and 0.4% respectively. Such conflicts occurred more often in SV than small variation alleles, and were more frequent in divergent assemblies for cactus (**Supplementary Figure 4**). VCF decomposition took 142- and 50-times longer for cactus and pggb compared to minigraph respectively, using approximately 26- and 22-times more memory (**Supplementary Table 1**), although pggb and cactus also contain substantial small variation to process.

The merging approach has an influence on the overlap of SVs between the three pangenomes. Merging based on variant breakpoints and length identified 221k nonredundant SVs ≥ 50 bp from the three pangenomes of which 135k (61.2%) were common (**Figure 2a**). Cactus contained 184k SVs of which 26k (14.1%) were private, i.e. not present in minigraph or pggb. We recovered fewer SVs with pggb (172k) and minigraph (169k) of which 8k (4.7%) and 9k (5.0%) were respectively private. The overlap between the three pangenomes was slightly higher (61.7%) for 112k SVs in non-repetitive regions, which span approximately 1.2 Gb (50.73%) of the ARS-UCD1.2 sequence. Consequently, the overlap was slightly lower (60.8%) for 108k SVs in repetitive regions. Applying stricter requirements to merge SVs across the pangenomes, requiring variants to be within a 5 bp window, had limited effect, and even requiring exact basepair resolution had 92k overlapping SVs, suggesting most SVs are precisely represented in the pangenomes (**Supplementary Figure 5**).

**Figure 2:**
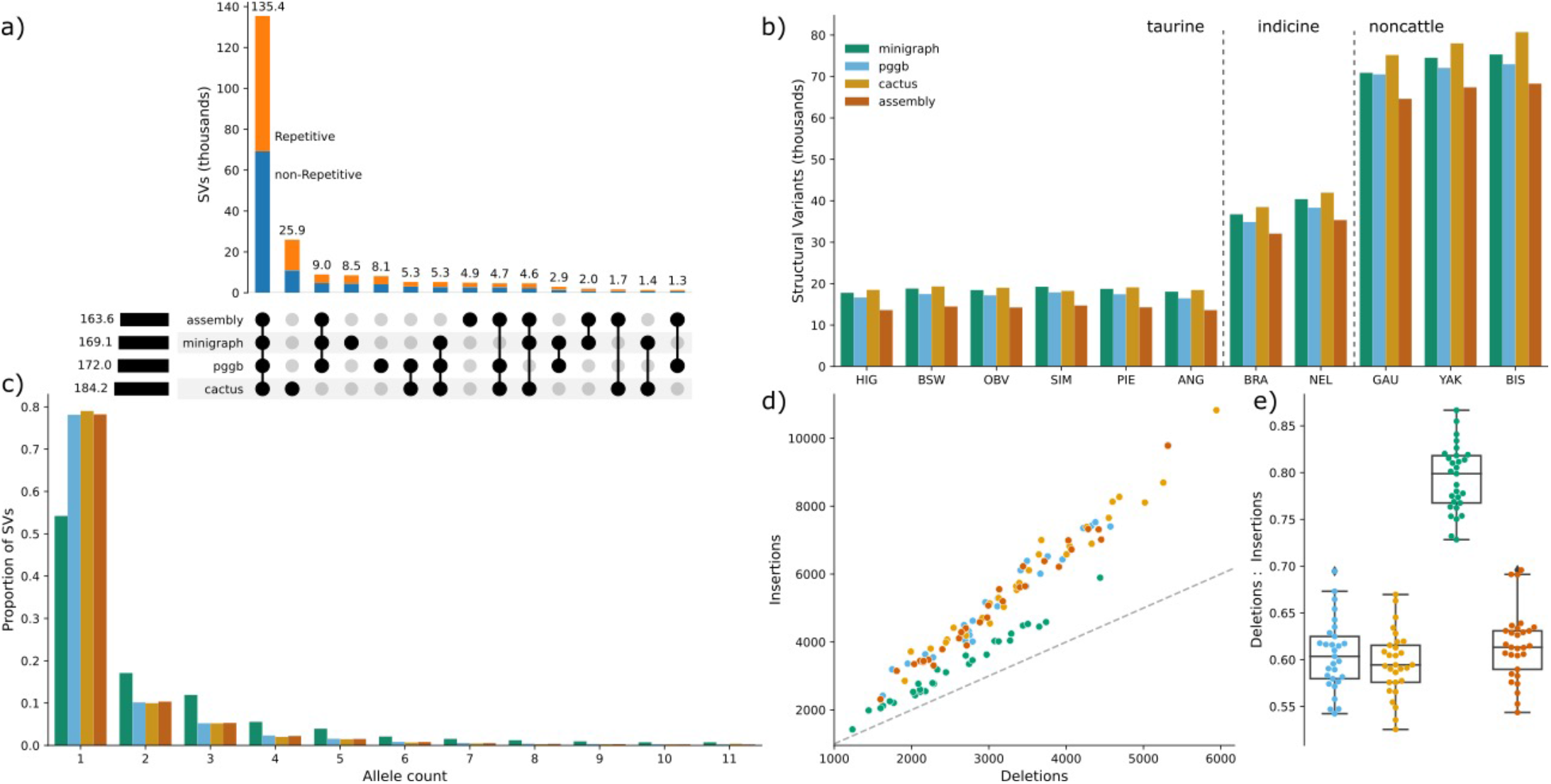
Consensus of structural variation between three pangenomes graphs. a) Overlap of SVs between three pangenomes and assembly-based SV discovery. Overall bar height represents the total number of SVs. Orange/blue colours represents SVs in repetitive/non-repetitive regions. b) Number of SVs identified in the input assemblies through assembly-based mapping and the three pangenomes. c) Allele count of the SVs. d) and e) Number of and ratio between deletions and insertions recovered from the pangenomes and assembly-based mapping.

A benchmark dataset to validate SVs is not available for Bovinae, and so we assessed the SV representation accuracy in the three pangenomes comparing against SVs called directly from reference-alignment with the same set of assemblies. We identified between 13.6k and 68.3k SVs in the haplotype-resolved assemblies, with substantially more SVs in gaur, yak, and bison than the indicine and taurine haplotypes (**Figure 2b**). More than three quarter (78.22%) of the SV alleles were observed in only one haplotype assembly. The eleven haplotype assemblies contained 163k nonredundant SVs of which 151k (92.30%), 150k (91.90%) and 146k (89.45%) were also recovered with minigraph, pggb and cactus, respectively. This approach validated the vast majority (N=135k, 96.26%) of the 141k SVs that were found in all three pangenomes. However, it validated only 6.12%, 14.06% and 19.24% of the SVs that were respectively private to cactus, pggb and minigraph. Similar alleles are collapsed more frequently in minigraph than pggb and cactus, and so the allele frequency spectrum differs between the three tools with minigraph containing substantially less singleton SVs (**Figure 2c**). While all tools recovered more insertions than deletions, minigraph contained proportionally less insertions than pggb and cactus (**Figure 2d,e**).

We used optical mapping from two unrelated Hereford and two unrelated Nellore samples [16] to further validate SV consensus in the pangenomes. The overlap is expected to be low due to the unrelated origin of samples, only covering two of the twelve breeds, and the different variant sizes accessible with optical mapping. Pggb had the largest number of validated SVs (1,661, 0.97%), followed closely by minigraph (1,656, 0.98%) and assembly (1,584, 0.97%) whereas cactus (1,218, 0.66%) had lower support (**Supplementary Figure 6**). Out of the pangenomes, minigraph had the most private SVs validated (37), compared to pggb (28) and cactus (6). We further assessed the commonality of small variation (SNPs and indels <50 bp) in pggb and cactus, as minigraph primarily only represents SVs. We recovered 68.27 M nonredundant small variants from the two pangenomes. Pggb and cactus contained 63.02 M (55.37 M SNPs / 7.65 M indels) and 64.27 M (56.61 M SNPs / 7.66 M indels) small variants of which 59.02 M (53.28 M SNPs / 5.74 M indels) were common. The overlap between pggb and cactus was substantially larger for SNPs than indels and multiallelic variants (**Figure 3a-c)**.

**Figure 3:**
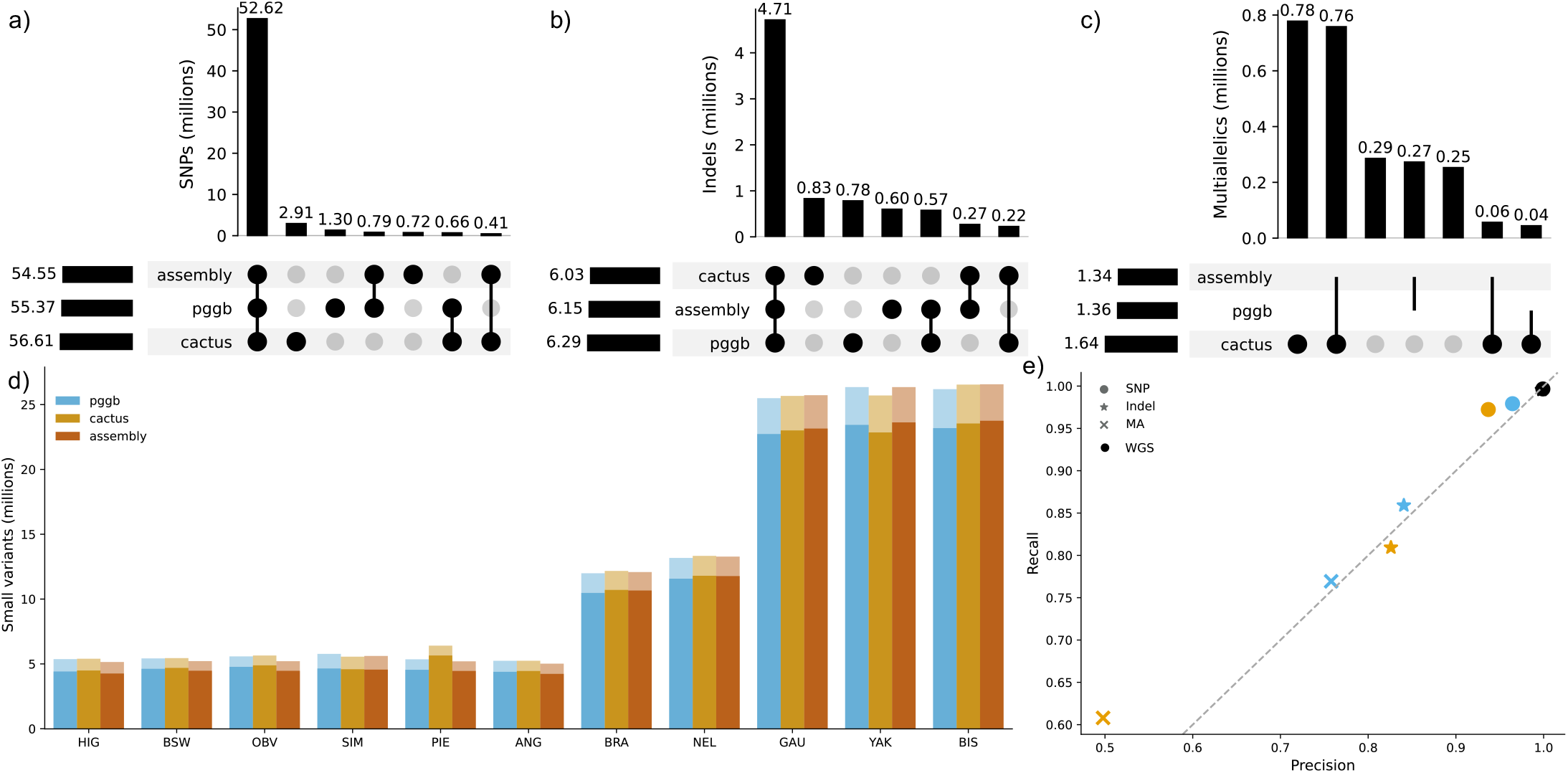
Overlap of SNPs and indels between pangenome- and assembly-based discovery. a-c) Overlap of small variations (SNPs and indels smaller then 50 bp) between pggb, cactus, and assembly-based discovery presented for (a) biallelic SNPs, (b) biallelic indels, and (c) multiallelic SNPs/indels. d) Number of small variations recovered from each haplotype from the pangenome and the assembly-based mapping. The faded and dark areas of the bars represent indels and SNPs, respectively. (e) Precision and recall of pggb and cactus for SNPs, indels, and multiallelic variants assuming the assembly-based calls are truth. The black WGS point represents gold-standard accuracy for 30x sequencing coverage.

We assessed small variation representation accuracy in the two pangenomes again using assembly-derived calls as an approximate truth set. The eleven input assemblies contained 62.04 M (54.55 M SNPs / 7.49 M indels) nonredundant small variations. Each assembly had between 5.12 and 27.05 M small variations, again finding substantially fewer in the taurine assemblies than their indicine and non-cattle counterparts (**Figure 3d**). Pggb and cactus contained 96.3% and 94.8% of the assembly-based small variants, and 94.8% and 91.5% of the pangenome variation respectively was found in the assembly calls (**Figure 3e**). Pggb had a higher overall F-score than cactus, 0.96 and 0.93 respectively, suggesting it encapsulates small variation more accurately.

### Alignment of assemblies to pangenomes

We realigned all twelve input assemblies (including ARS-UCD1.2) against the pangenomes to calculate edit distances and quantify sequence content of the pangenomes. Since centromeric, and to a lesser extent telomeric, sequence is challenging to align, we analysed these “centro-/telomeric” regions separately to the rest of the “bulk” sequence (**Supplementary Table 2**). Minigraph had a substantially higher edit rate (0.494%) than pggb (0.010%) and cactus (0.010%) in bulk regions and also in centro-/telomeric regions, 3.43%, 0.396%, and 0.719% respectively (**Figure 4a**). Even though cactus includes more centromeric sequence in the graph (**Supplementary Figure 1**) and had overall comparable edit rates, its centro-/telomeric edit rates were nearly double that of pggb. Minigraph also had a significantly lower edit rate for ARS-UCD1.2, as all reference sequence is included by definition of minigraph’s algorithm.

**Figure 4:**
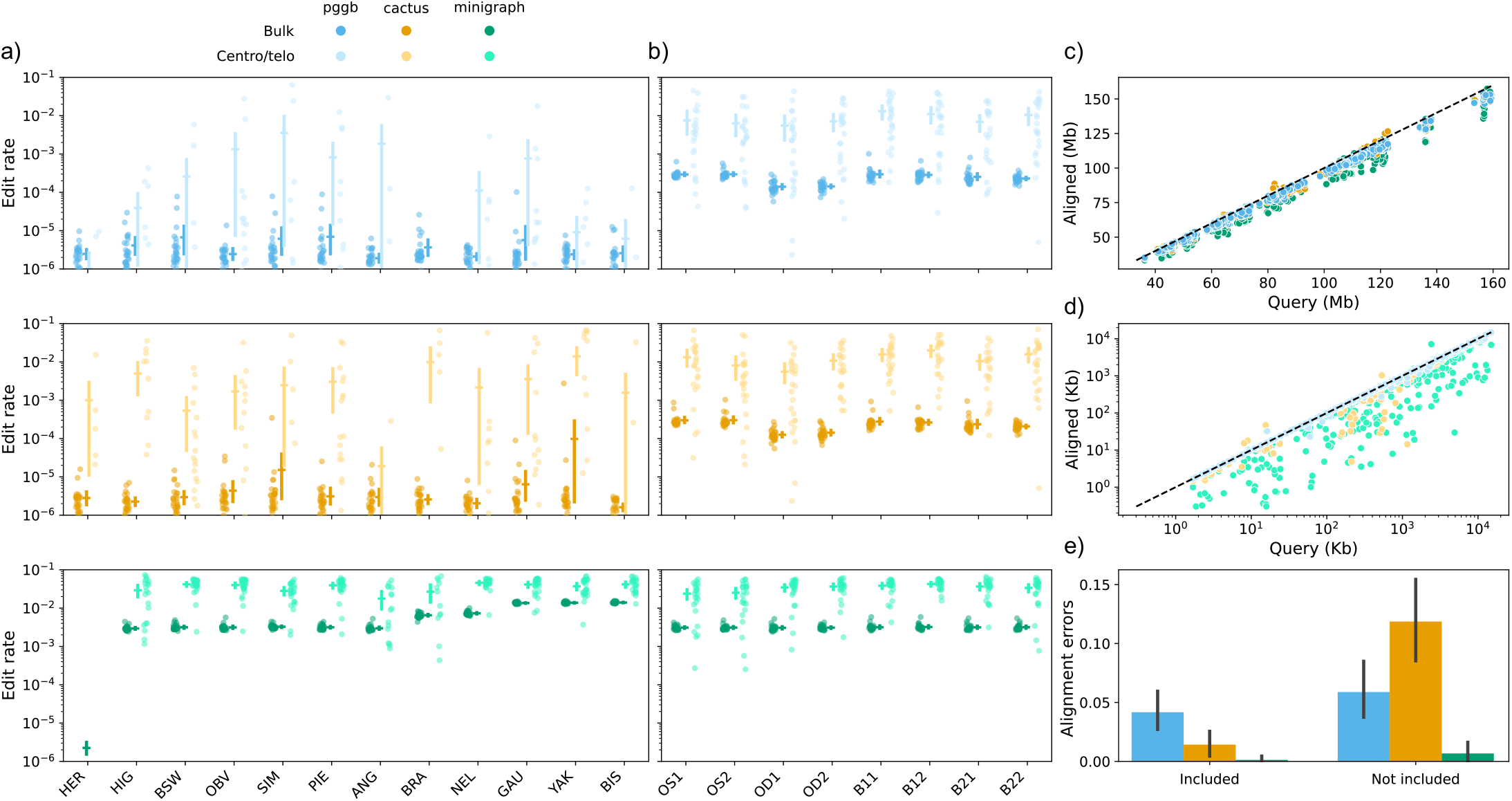
Realignment of included and held-out assemblies to the pangenomes. a) Edit rate of the twelve assemblies used in pangenome construction. Each faded dot represents one autosome of each assembly in either the bulk or centro-/telomeric ranges. Mean and 95% confidence intervals are also indicated for each category of sequence per assembly. b) Similar plot to (a) but for eight additional assemblies held-out from pangenome construction. c) Query coverage for the bulk sequence for each autosome of all assembly. Values above the dashed line indicate there were more aligned bases than query bases, suggesting multiple ambiguous alignments that were equally scored. d) Similar to (c), but for the centro-/telomeric sequence. e) Bar plots of for the number of alignment errors reported by GraphAligner when using finite “tangle effort”. Bar heights reflect the mean across 348 (Included) and 232 (Not included) autosomes across the assemblies, while error bars represent 95% confidence intervals.

Edit rates were higher for assemblies more diverged from the reference backbone in minigraph pangenomes, indicating they do not incorporate highly diverged segments containing multiple small variations but were not large enough to form bubbles. Contrastingly, pggb and cactus pangenomes did not have any divergence bias for edit rate, suggesting they may be more suited to include more divergent assemblies into “super-pangenomes” [17]. However, there was a weak increase in edit rate in pggb and cactus pangenomes for HiFi assemblies, potentially related to the increased resolution of complex and repetitive areas beyond centro- /telomeric sequence that cannot be realigned accurately.

We also aligned eight additional assemblies held out from the pangenomes (2x haplotypes from each parent of the OBV and 2x haplotypes for two unrelated Brown Swiss cattle [**Figure 1**]), and found highly similar edit rates for minigraph, but significantly higher edit rates for pggb and cactus (**Figure 4b**). This is expected and reflects true genomic variation in the held-out assemblies not present in the pggb/cactus pangenomes, while minigraph never captured small variation anyway and had higher edit rates to begin with. The OD1/2 (dam of OBV sample) assemblies had a lower edit rate in pggb and cactus, as expected since the OBV haplotype is the maternal haplotype.

In addition to substantially lower edit rates, pggb and cactus also covered more of the bulk query sequence (98.4% and 98.6% respectively) compared to minigraph (93.8%). The differences were more pronounced in the centro/telomeric query sequence alignments (98.0%, 93.9%, and 65.8% respectively), as expected given the relative lack of centromeric sequence in minigraph pangenomes (**Figure 4c**). Some chromosomes (e.g. 6, 14, and 16) had multiple ambiguous but equally scored alignments in cactus pangenomes, leading to >100% query coverage. More query sequence can be aligned with more sensitive alignment parameters, but this requires >300 GB of memory per chromosome in cactus graphs (S**upplementary Table 3**). Even with relaxed alignment parameters, aligning to cactus and pggb pangenomes took 2.2- and 1.4-times more CPU time and 37- and 7-times more memory than to minigraph pangenomes (**Supplementary Table 4**), and was especially prounced for aligning assemblies not included in the pangenomes. Failed alignment of 500 Kb segments were more common in pggb and cactus than minigraph, generally resulting from exceeding GraphAligner’s “tangle effort” in highly complex regions (**Figure 4e**). This was especially apparent when aligning not included assemblies to the cactus pangenomes.

### Pangenome resolution of VNTR

Variable number tandem repeats (VNTR) account for substantial gene expression and complex trait variation, but due to their repetitive nature and high mutation rates these loci are difficult to resolve from linear alignments [18],[19]. A catalogue of bovine VNTR had not been established and so we identified 9,568 tandem repeats (TRs) in non-masked regions of the ARS-UCD1.2 reference sequence (**Supplementary Table 5**) and investigated their prevalence in the other assemblies through the three pangenomes.

Pggb and cactus contain all assembly sequence, and so can arbitrarily convert between pangenome position and assembly coordinate with tools like odgi. As such, we can liftover the TR positions directly into each assembly through the pangenomes. On the other hand, minigraph contains information on assembly coordinates at graph bubbles, and so we can only estimate TR positions that effectively overlap with SVs. The former approach is substantially more complete and accurate, but incurs a much larger compute cost (**Table 2**). Approximately 95% of TRs had all twelve assemblies associated with a pangenome path, while the remaining TRs suffered from reduced mappability (**Table 2**). All pangenomes had instances where coordinates were erroneously translated, but this was most apparent in cactus with several huge outliers (**Supplementary Table 6**). Nearly all TRs examined by minigraph were variable between the assemblies, while approximately 12% of TRs examined by pggb and cactus had zero variation between samples (**Table 2**).

**Table 2:** All TRs were examined from pggb and cactus, while TRs overlapping SVs were examined for minigraph. Genotypable TRs are TRs where all 12 assemblies had a known path through the local graph structure. VNTRs are genotypable TRs that had at least one sample with a different number of TR counts. CPU and Memory indicate the compute resources needed to identify the paths of all assemblies through all examined TRs.

| Pangenome | TRs examined | Genotypable TRs | VNTRs | CPU (h) | Memory (Gb) |
| --- | --- | --- | --- | --- | --- |
| Pggg | 9566 | 8939 (93.5%) | 7854 (87.9%) | 18.17 | 7.5 |
| Cactus | 9566 | 8910 (93.1%) | 7859 (88.2%) | 24.17 | 7.7 |
| Minigraph | 5731 | 5504 (96.5%) | 5457 (99.1%) | 0.19 | 0.5 |

Tandem repeat counts were similar between the three pangenomes, with 5,293 commonly genotyped VNTRs. There were 3,332 (63%) VNTRs with identical counts across all 12 assemblies in all three pangenomes and 4,084 (77%) identical in at least two pangenomes (**Figure 5a**). The remaining VNTR counts still broadly agreed with minor variability (**Supplementary Figure 7**). The average squared Spearman correlation was 0.92 between the pangenomes, with several outliers in cactus and minigraph skewing the squared Pearson correlation, which was otherwise around an average of 0.8 (**Figure 5b**, all significant). TRs present in only pggb and cactus had low variability between cattle and non-cattle (**Figure 5c**), suggesting minigraph efficiently captures nearly all VNTRs of interest for further investigation. We also genotyped 465 TRs with adVNTR [19] using HiFi reads from the gaur, Nellore, Piedmontese, Brown Swiss, and Original Braunvieh samples as an approximate truth set, finding good concordance (**Figure 5d, Supplementary Table 5**), with median difference of counts of 7, 8, and 11 for pggb, cactus, and minigraph to adVNTR. Minigraph occasionally over- or underestimated TR counts if the overlapping SV was significantly larger or smaller, as that determines the translated coordinates. There were also instances where all three pangenomes agreed on counts different to adVNTR, indicating even HiFi reads may not be powerful enough to genotype all VNTRs (**Figure 5e**). Clustering based on TR counts produced trees that broadly grouped the taurine and indicine cattle as well as the non-cattle (**Figure 5f**), although the minigraph and cactus trees had slight inconsistencies with the topology of the mash-derived tree (**Figure 1d**).

**Figure 5:**
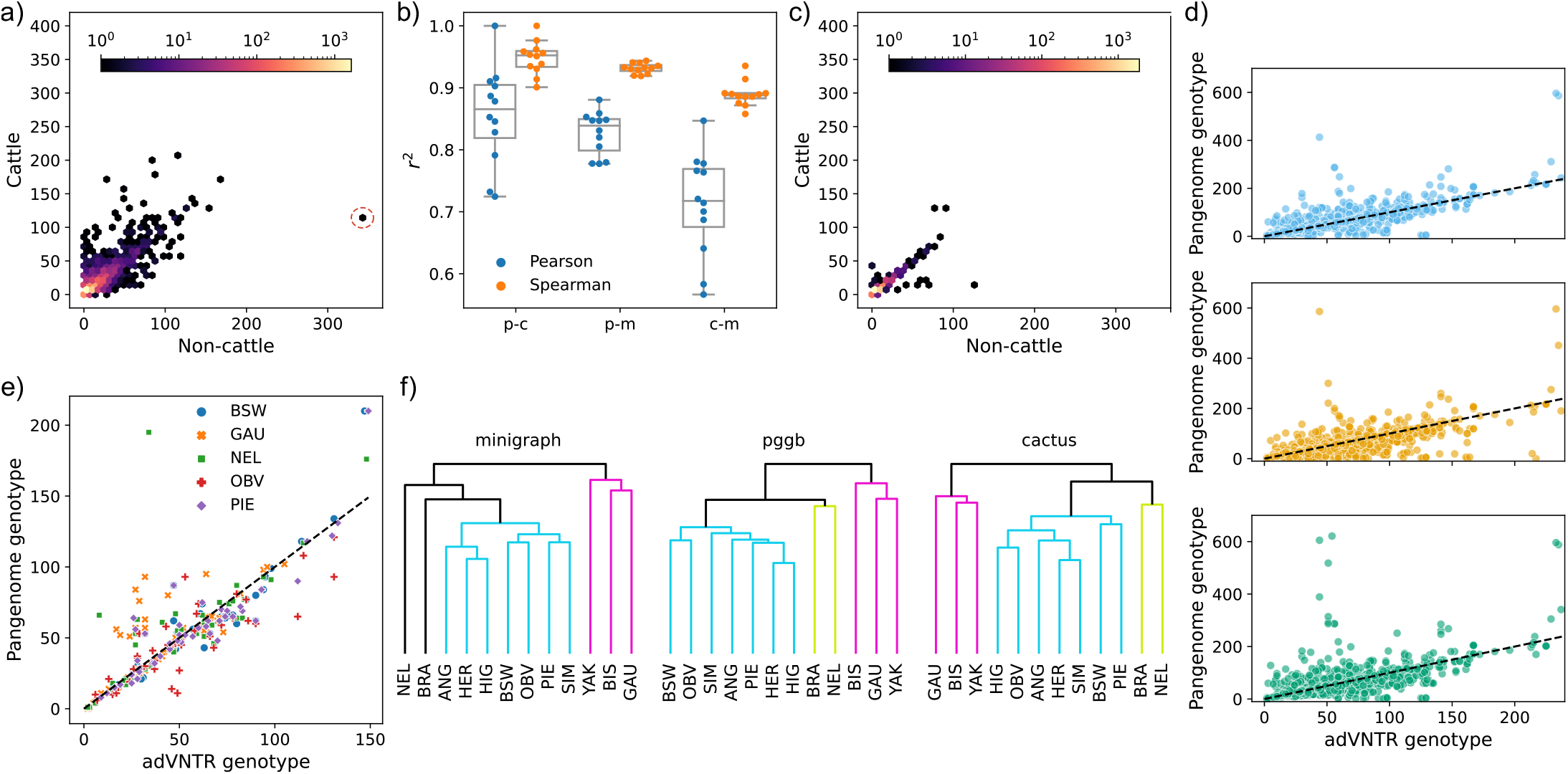
VNTR concordance in three pangenomes. a) TRs with identical counts in at least two pangenomes, with the median count for the cattle and non-cattle groups. A particular VNTR with substantially more repeats in non-cattle compared to cattle that we investigated further is circled in red. b) Pearson and Spearman squared correlation coefficients across the TR counts for pggb-cactus (p-c), pggb-minigraph (p-m), and cactus-minigraph (c-m). Each point is one assembly, with box plots over the 12 assemblies. c) Similar to (a), except TRs with identical counts in pggb and cactus that were not present in minigraph. d) adVNTR-derived genotypes for five HiFi samples in the three pangenomes. The black dashed line indicates the expected count using adVNTR as a ground truth. e) VNTRs where all three pangenomes agreed to a different count than adVNTR, suggesting adVNTR may sometimes over-/underestimate assembly-based counts. f) Trees derived from the TR counts across different input assemblies, with colours representing clusters of taurine cattle, indicine cattle, and non-cattle.

### An eVNTR mediates expression of neighboring genes and non-coding RNA

A VNTR locus on chromosome 12 (86,161,072 - 86,162,351 bp) was highly variable, containing substantially more copies of a degenerate 12 bp motif in non-cattle and fewer in Nellore than in the taurine assemblies (**Figure 6a,b**), prompting a detailed investigation. Viewing this VNTR in a Bandage plot revealed different structure in the three pangenomes, as well as the motif variation present (**Figure 6c, Supplementary Figure 8**).

**Figure 6:**
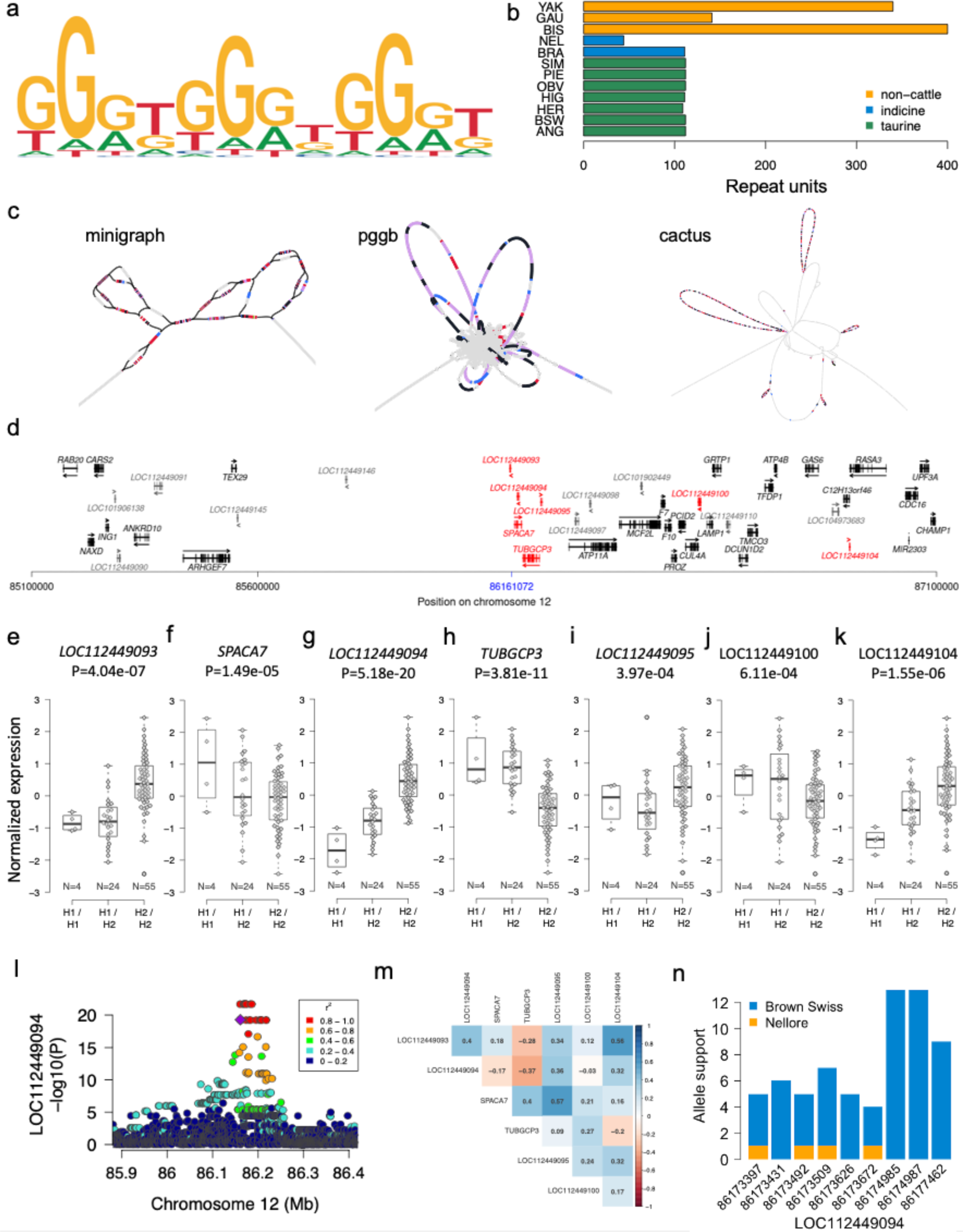
A polymorphic eVNTR discovered from the pangenomes. a) VNTR repeat motif and b) number of repeat units in the twelve input assemblies. c) Bandage plots of a VNTR upstream *SPACA7*. BLAST hits for the top 5 most common VNTR motifs are coloured per motif. d) Genes (black) and (long) non-coding RNAs (dark grey) nearby the VNTR. Blue colour indicates the position of the VNTR. Arrows indicate the orientation of genes and lncRNA. Red colour indicates two genes and five lncRNAs whose expression is associated with the VNTR. e-k) Expression (quantified in transcripts per million (TPM)) of the associated genes and transcripts in testis tissues of 83 Brown Swiss bulls that are either homozygous for hap1 (H1 / H1), homozygous for hap2 (H2 / H2) or heterozygous (H1 / H2). The number of animals per diplotype is below the boxplots. l) Cis-expression QTL mapping for *LOC112449094*. Different colours indicate the pairwise linkage disequilibrium (r^2^) between the VNTR haplotype (violet) and all other variants. m) Correlation between the abundance of mRNA and lncRNA for the associated genes and lncRNA. n) Allelic imbalance of nine heterozygous exonic SNPs in *LOC112449094* in testis tissue of a Nellore x Brown Swiss crossbred bull. Orange and blue colours represent paternal (Nellore) and maternal alleles (Brown Swiss).

We observed two VNTR “haplotypes” (referred to as hap1 and hap2) in long read alignments of additional Original Braunvieh cattle demonstrating within-breed variability. While the alignments indicated several insertions and deletions in both haplotypes with respect to ARS-UCD1.2, hap2 is 24 bp longer than hap1 due to two additional copies of the repeat motif (**Supplementary Figure 9**). There are 63 genes annotated within the *cis*-regulatory range (±1 Mb) of the VNTR, of which 44 (**Figure 6d**) were expressed at >0.2 TPM in testis tissues of 83 mature Brown Swiss and Original Braunvieh bulls.

A degenerate repeat motif and an overall length >1300 bp precluded the short sequencing read-based genotyping of the VNTR in the eQTL cohort with adVNTR. Instead, we genotyped the VNTR through two SNPs (Chr12:86160984 and Chr12:86160971) and one indel (Chr12:86161000) that tagged the two VNTR haplotypes. This enabled us to investigate putative *cis*- and *trans*-regulatory impacts of the VNTR on the expression of genes and long non-coding RNAs (lncRNA). Two genes (*TUBGCP3, SPACA7*) and five non-coding RNAs (*LOC112449093, LOC112449094, LOC112449095, LOC112449100, LOC112449104*) within ±1 Mb of the VNTR were differentially expressed (P<1.13e-3, Bonferroni-corrected significance threshold) between the diplotypes indicating a putative *cis*-regulatory role (**Figure 6e-k**). We did not detect *trans*-regulatory effects for the haplotype (**Supplementary Figure 10**).

We mapped eQTL within the *cis*-regulatory range (±1 Mb of the transcription start site) of the two significant genes and five significant non-coding RNAs to investigate if variants in *cis* other than the VNTR haplotype are associated with transcript abundance. The VNTR haplotype was among the top variants at the *LOC112449094* eQTL, but eleven variants in strong linkage disequilibrium (r^2^= 0.93) were more significant (P=2.00e-22 *vs*. P=5.18e-20) (**Figure 6l**). The eQTL peak was absent when the VNTR haplotype was fitted as a covariate. We made similar observations for the *TUBGCP3* eQTL (**Supplementary Figure 11**). Variants other than the VNTR haplotype were more strongly associated with the expression of *SPACA7, LOC112449093, LOC112449095, LOC112449100*, and *LOC112449104* (**Supplementary Figure 11**). These findings suggest that the VNTR haplotype primarily mediates the expression of *LOC112449094* and *TUBGCP3*.

Hap2 containing more copies of the 12 bp motif increases abundance (β_normalized_=1.26 ± 0.10, P=5.18e-20) of *LOC112449094* which is lowly-expressed (0.86 ± 0.31 TPM) in 83 bull testis transcriptomes. Conversely, hap2 is associated (β_normalized_= -1.08 ± 0.14, P=3.81e-11) with lower *TUBGCP3* mRNA. Negative correlation (–0.37, **Figure 6m**) between *LOC112449094* and *TUBGCP3* abundance suggests that *LOC112449094* could be a cis-acting lncRNA that represses expression of *TUBGCP3*. The VNTR haplotype is in strong LD with two SNPs (Chr12:86173431 (r^2^=0.87) and Chr12:86173509 (r^2^=0.93)) in *LOC112449094* exons. While hap2 primarily segregates with the alternate alleles of these two SNPs, hap1 segregates with the respective reference alleles. Alternate allele support in RNA sequencing reads overlapping Chr12:86173431 and Chr12:86173509, respectively, was 71% (96 out of 135 alleles, P_binom_=1.01e-06) and 69% (94 out of 137 alleles, P_binom_=1.57e-05) in 22 animals that are heterozygous both at the VNTR haplotype and the SNPs confirming allelic imbalance due to lower *LOC112449094* expression in hap1.

We confirmed a putative *cis*-regulatory role of the VNTR in an individual with indicine ancestry. Expression of *LOC112449094* was relatively low (TPM=0.66) in testis tissue of a Nellore (*Bos taurus indicus*) x Brown Swiss (*Bos taurus taurus*) crossbred bull (NxB), although it inherited hap2 from its Brown Swiss dam (**Supplementary Figure 9**). The paternal (Nellore) haplotype is diverged from hap2 and hap1, and contains less VNTR repeat units (F**igure 6b**). Alignment of parental-binned HiFi reads against ARS-UCD1.2 readily distinguished paternal from maternal alleles at nine heterozygous SNPs overlapping *LOC112449094* exons. Inspection of these variants in the RNA sequencing alignments of NxB F1 showed allelic imbalance due to a disproportionally low number or even complete absence of paternal (=Nellore) alleles (**Figure 6n**) suggesting that the low VNTR repeat count of the Nellore haplotype represses expression of *LOC112449094*.

## Discussion

We constructed pangenomes using three different algorithmic approaches with minigraph, pggb, and cactus, finding each have different strengths and weaknesses. Pggb and cactus include all variation, down to SNPs, while minigraph primarily captures SVs. Consequently, pggb and cactus pangenomes had roughly 450 times more nodes and edges, 10 times the file size, and construction required 253 and 16 times as much CPU time as the minigraph pangenomes respectively, despite only including about 5 times as much non-reference sequence. High-quality assemblies are becoming increasingly easier and cheaper to produce, and so expanding pangenomes with additional assemblies will likely be common. Minigraph can iteratively add assemblies in near-linear time and memory to an existing pangenome. Cactus requires storage of modestly sized intermediate files (10s of GB) and the complexity of adding a new assembly depends on the existing phylogenetic guide tree [5]. Pggb on the other hand, performs all-versus-all alignment, and so adding new assemblies would require building a new pangenome from scratch. As such, pggb may be suited to building reference pangenomes or periodic updates, but cactus and minigraph are more appropriate for projects with ongoing assembly efforts.

All three pangenomes contained more SVs than the assembly-based “truth”, with many private SVs in minigraph (8.5k), pggb (8.1k), and cactus (25.9k), likely representing false positives, agreeing with recent findings in human pangenomes [13]. However, the overall SV agreement was good, with minigraph, pggb, and cactus respectively having F-scores of 0.91, 0.90, and 0.84. Minigraph and pggb furthermore had the best overlap with orthogonal optical mapping data, supporting many of their SVs are true variation. We found that approximately 0.5% and 3.9% of alleles in pggb and cactus were removed during preliminary filtering when only keeping alleles seen in genotypes, suggesting variants may either exist in complex regions that cannot be genotyped confidently or strongly tangled regions that produce spurious variant calls. However, we found a higher overlap on small variations compared to structural variations, indicating that the cactus (F1: 0.93) and pggb (F1: 0.96) pangenomes can integrate small variations more accurately. These results demonstrate that SVs and small variation encoded in pangenome graphs, particularly cactus, would benefit from careful filtering before qualifying for downstream use in population genomic analyses.

Pggb and cactus are reference-free, while minigraph requires an initial backbone to determine variation from the input assemblies. Minigraph is thus susceptible to reference-bias, especially in the case of an incomplete reference (currently almost every reference genome except the T2T-CHM13 [9]). The ARS-UCD1.2 autosomes are interrupted by 257 gaps and contain almost no centro-/telomeric sequence, which prevents integration of centro-/telomeric sequence from the more complete HiFi assemblies into the minigraph pangenome. Pggb and cactus are largely able to integrate these sequences, with 255 and 292 Mb of centromeric sequence in the non-reference nodes of the pangenomes, respectively. However, cactus does use progressive alignments on the provided guide tree, and so does appear to have a slight reference-bias when projecting back into a reference coordinate system (like VCF). Assemblies more distant to the ARS-UCD1.2 branch tended to have higher genotype conflict rates, which is not observed in pggb.

Pangenomes with base-level variation like pggb and cactus can realign most of the input assemblies with minimal edit rate, but centro-/telomeric regions are still poorly realigned. Equally, the base-level variation in these pangenomes can lead to strongly tangled graph topology, which can generate poor realignments and substantially increase computational requirements. For super-pangenomes incorporating multiple (sub-)species, containing a greater amount of variation, downstream software like Bandage, GraphAligner, and vg quickly hit CPU and memory bottlenecks working with pggb or cactus, while minigraph pangenomes are usable even without HPC infrastructure. Although beyond its design intentions, minigraph can be run with a lower “minimum variant length” parameter, increasing its sensitivity to smaller variation down to approximately 10 bp without substantially impacting compute requirements for pangenome construction or analyses (**Supplementary Table 7**). However, this unsupported use-case cannot regularly capture SNPs and so pggb and cactus are vastly better for representing small variation accurately.

We show that the three pangenomes are generally concordant at highly polymorphic VNTR loci, although minigraph can over- or underestimate TR counts if the graph bubbles are significantly larger or smaller than the TR region, as minigraph primarily translates assembly coordinates at these bubbles. All three pangenomes had several poorly genotyped VNTRs, due to incorrect coordinate extraction in these repetitive regions, again indicating unresolved challenges with complex alignments. Furthermore, minigraph primarily captures repeat count variation rather than motif variation in TRs, the latter of which may be of greater importance in identifying eVNTRs [20]. However, the small variation present in pggb and cactus hinders visual inspection and pangenome annotation with Bandage, while TRs are simple to identify in SV-oriented minigraph pangenomes. We also show that even highly accurate long reads can fail to successfully genotype VNTRs, supporting that assembly-based approaches are the most powerful resource for detecting large or complex variation. In the case of a highly variable eVNTR upstream *SPACA7*, adVNTR predicted 116 copies for gaur, whereas both pggb and minigraph pangenomes identified 141 copies. Cactus erroneously aligned several assemblies in this VNTR region, and so only predicted 53 copies for gaur but correctly predicted the taurine cattle count.

## Conclusions

Bovine pangenomes have been utilized before to make non-reference sequences amenable to association testing and reveal trait-associated structural variants [4],[15],[7]. Our findings show good agreement between the SVs discovered from three widely used pangenome methods and so reinforce the utility of any pangenome approach to investigate variants that are difficult to resolve with a single linear reference. Pggb, and to a lesser extent cactus, pangenomes losslessly represent the input assemblies and are ideal for generating a pangenome reference containing all variation. However, minigraph has much greater utility, allowing simple expansion with additional assemblies and rapid downstream analyses with modest computational requirements. With the recent establishment of the cross-species Bovine Pangenome Consortium (https://bovinepangenome.github.io/) and similar efforts in plants [17],[21],[22], there is a need to emphasise pangenomes that can handle significantly higher levels of variation compared to the draft human pangenome reference [13].

## Methods

### Pangenome construction

Pangenomes were constructed per-chromosome for the 29 bovine autosomes, using the Hereford-based *Bos taurus taurus* ARS-UCD1.2 reference genome [23] and eleven other haplotype-resolved assemblies: eight domestic cattle (*Bos taurus taurus* and *Bos taurus indicus*) and three wild relatives of domestic cattle (*Bos grunniens, Bos gaurus, Bison bison*) (**Table 3**). We used these assemblies to construct pangenomes with minigraph (v0.20) [3], the PanGenome Graph Builder (pggb, v0.5.2) [24], and the Cactus Progressive pangenome pipeline (cactus v2.3.0) [5].

**Table 3:** Input assemblies for the pangenomes. The bovine reference (Hereford, ARS-UCD1.2) is a primary assembly, a collapsed haploid representation of the diploid genome. All other assemblies are haplotype-resolved, a haploid genome of either the maternal or paternal haplotype. CLR: Pacific Biosciences Continuous Long Reads, HiFi: Pacific Biosciences High Fidelity Reads, ONT: Oxford Nanopore Technologies Reads.

| Breed/species | Acronym | Primary sequencing strategy | Autosome length (Gb) | Reference |
| --- | --- | --- | --- | --- |
| Hereford / *Bos taurus taurus* | UCD | CLR | 2.489 | [23] |
| Angus / *Bos taurus taurus* | ANG | CLR | 2.468 | [14] |
| Brahman / *Bos taurus indicus* | BRA | CLR | 2.478 | [14] |
| Highland / *Bos taurus taurus* | HIG | ONT | 2.483 | [25] |
| Yak / *Bos grunniens* | YAK | ONT | 2.478 | [25] |
| Simmental / *Bos taurus taurus* | SIM | ONT | 2.494 | [26] |
| Bison / *Bison bison bison* | BIS | ONT | 2.488 | [27] |
| Brown Swiss / *Bos taurus taurus* | BSW | HiFi | 2.617 | [7] |
| Nellore / *Bos taurus indicus* | NEL | HiFi | 2.602 | [7] |
| Piedmontese / *Bos taurus taurus* | PIE | HiFi | 2.560 | [7] |
| Gaur / *Bos gaurus* | GAU | HiFi | 2.520 | [7] |
| Original Braunvieh / *Bos taurus taurus* | OBV | HiFi | 2.532 | [7] |

The minigraph pangenome was constructed using base-level alignment (‘-c‘), 2% divergence level, and default parameters otherwise. The ARS-UCD1.2 reference genome was the backbone of the graph, and the other assemblies were added in order of their mash distance [28] to the reference: Highland, Brown Swiss, Original Braunvieh, Simmental, Piedmontese, Angus, Brahman, Nellore, gaur, yak, and bison.

Pggb was run with the recommended parameters (segment length of 100000 bp, identity mapping taken from the rounded lowest mash similarity of 98%, and target haplotype paths of 12) with all assemblies combined into a single fasta file.

Since our assemblies contain multiple species with significant divergence, we used cactus with progressive alignment using the guide tree from the mash-based distance. Assemblies were first soft-masked with the rush job mode of RepeatMasker (http://www.repeatmasker.org) (version 4.1.4) and rmblast (version 2.13.0), using a database of repetitive DNA elements from Repbase (release 20181026). The cactus hierarchical alignment (HAL) was converted into GFA using hal2vg (https://github.com/ComparativeGenomicsToolkit/hal2vg) and vg convert [29].

### Analysis of pangenome sequence content

We extracted all fasta sequence in the graphs with odgi flatten (v0.8) [24], and then repeat masked the output as described above.

### Variation discovery from pangenomes

Pangenomes were decomposed into VCF files using vg deconstruct with --all-snarls --path-traversals –ploidy 1. We added path information (P-lines) to minigraph’s gfa by manually curating the output of minigraph -- call on each sample which retraces the assembly’s path (available in the Github repository associated with this paper). Variants were then normalized using vcfwave [31], skipping variants larger than 1 Mb, and then split into small (<50 bp) and structural variation (>50 bp) using bcftools norm and bcftools view. Alleles not observed in any sample were dropped using --trim-alt-alleles during splitting, and similarly applied to per-sample VCF statistics. Multiple nucleotide polymorphisms (MNPs) were split into multiple SNPs using bcftools norm --atomize.

### Variation discovery from assemblies

All non-reference assemblies were aligned to ARS-UCD1.2 using minimap2 (v2.24 [32]) with “-cx <preset>“. The divergence-based preset was chosen as asm5, asm10, and asm20 for the taurine, indicine, and non-cattle, respectively. Variants were called from the alignments using paftools.js and normalized and subsetted using the same approach as pangenome variation.

### Variation consensus

SVs from the three pangenomes and assemblies were merged using Jasmine (v.1.1.5 [33]) with -- normalize_type --allow_intrasample max_dist=1000 max_dist_linear=0.5. A more lenient SV merging was done using parameters max_dist=10000 max_dist_linear=1.0. A more strict merging was done with max_dist=5 min_seq_id=0.5, and a near-perfect merging was done with max_dist=1 max_dist_linear=0 min_seq_id=0.85.

Jasmine was also used for intersection SVs with optical mapping data, using the lenient parameters described above. The optical mapping data filtered to keep deletions, insertions, and duplications and removing SVs >1 Mb.

Small variation between individual haplotypes was intersected using bcftools isec, requiring exact matches of REF and ALT alleles with the --none flag. The normalized cohort-level pangenome and assembly VCF files were intersected based on exact matches of small variant coordinates and ALT alleles.

### Calculation of edit distance

The twelve assemblies included in the pangenomes and eight haplotype assemblies from four taurine cattle which are not part of the pangenomes were realigned to the pangenomes using GraphAligner (v1.0.16, [34]) with parameters -x dbg -C 100000 --max-trace-count 5 --seeds-minimizer-ignore-frequent 0.001 --precise-clipping 0.9. Centro-/telomeric sequence intervals, as classified by RepeatMasker, were merged within 250 Kb and 1 Kb by bedtools [35] respectively, and then split into centro-/telomeric sequence and bulk sequence fasta files. Both files were split every 500 Kb during realignment for computational constraints. Edit rate and query coverage was then calculated from the resulting graph alignment files.

The eight additional haplotype assemblies were constructed with the dual assembly approach in hifiasm (v0.16.1, [36]) using between 17 and 24–fold HiFi read coverage from two purebred Brown Swiss and two purebred Original Braunvieh cattle. The two Original Braunvieh cattle are the sire and dam (labelled as OS1, OS2, OD1, OD2) of an OxO F1 which was the source for the OBV haplotype [7] included in the pangenomes whereas the two Brown Swiss cattle are not directly related to the BSW haplotype included in the pangenomes (labelled as B11, B12, B21, B22).

### VNTR analysis

We identified TRs in the ARS-UCD1.2 reference sequence using TRF ([37], v4.10.0) with parameters “2 7 7 80 10 50 500”, allowing a minimum and maximum motif length of 10 and 100 bp respectively and a minimum and maximum tandem repeat length of 50 bp and 10 Kb. We excluded TRs which overlapped with repeat elements identified in the reference sequence using bedtools intersect -v -f 0.5. For minigraph, we then used bedtools intersect again for the TRs and the minigraph SV bed files generated by gfatools bubble. We then extracted the approximate coordinates using minigraph --call for each assembly. For pggb and cactus we used the position command of odgi to translate the TR coordinates from ARS-UCD1.2 to the paths of other assemblies in the graphs.

We calculated the number of TRs by extracting per-assembly sequence from the VNTR coordinates predicted from the pangenomes. We then used the python regex package to count how often the TR repeat motif occurs in the query sequence, allowing 25% error rate to account for substitutions, insertions, and/or deletions between repeat units.

We used adVNTR (v1.4.1, [19]) to genotype the 465 VNTRs with at least 50 repeat units in one of our HiFi samples. We built a database for these TRs using adVNTR addmodel, and then genotyped them using adVNTR genotype with the flags “--naive --pacbio --haploid” on the triobinned HiFi samples. These bam files were generated using sequences binned by Canu (v2.2, [38]) using parental reads, followed by alignment to ARS-UCD1.2 with minimap2 (as described in **Variation discovery from assemblies** except using the map-hifi preset).

The VNTR annotated pangenomes were generated using Bandage [39], using its built-in BLAST feature. The blastDB is created from the graph sequence, while we provide common TR motifs for given VNTRs as queries. Since motifs are “short” compared to typical blast queries, we use “-task blastn-short -word_size <TR length> -evalue 100” and filter minimum alignment sizes below <TR length> to increase sensitivity to specific motifs.

### Establishing a testis eQTL cohort

Testis tissue of 83 mature bulls was collected at a commercial slaughterhouse and subjected to DNA and RNA purification as described earlier [40]. DNA samples were sequenced on an Illumina NovaSeq6000 instrument using 150 bp paired-end sequencing libraries. Following quality control, between 74,371,404 and 304,199,764 filtered read pairs per sample (mean: 268,798,344 ± 80,134,660) were aligned to the ARS-UCD1.2 reference sequence and processed as described in Kadri et al. [40]. Single nucleotide and short insertion and deletion polymorphisms were discovered and genotyped using DeepVariant (version 1.3.0, [41]). Beagle4.1 [42] was applied to impute sporadically missing genotypes and infer haplotypes. A genomic relationship matrix was constructed and subjected to a principal components analysis using plink (v.1.9, [43]).

Total RNA sequencing libraries (2 × 150 bp) were prepared using the Illumina TruSeq Stranded Total RNA sequencing kit and sequenced on an Illumina NovaSeq6000. Following quality control, between 70,249,355 and 177,053,828 filtered read pairs per sample (mean: 130,092,907 ± 18,684,899) were aligned to the ARS-UCD1.2 reference sequence and the Refseq gene annotation (release 106) using the splice-aware read alignment tool STAR (version 2.7.9a) [44] with options --twopassMode Basic and --waspOutputMode SAMtag to enable robust and unbiased allele-specific expression detection [45]. Per-individual VCF files required for WASP filtering were prepared from the cohort-level VCF files (see above).

Testis tissue from the NxB F1 was collected after regular slaughter. Total RNA was prepared and sequenced as described above. Following quality control, 136,820,136 filtered read pairs were aligned against the bovine reference sequence using STAR (see above).

### Gene expression quantification

The expression level of genes, ncRNA and lncRNA was quantified (in transcripts per million, TPM) using kallisto (version 0.46.1, [46]) and aggregated to the gene level using the R package tximport [47]. We retained genes, ncRNA and lncRNA that were expressed at an average value >0.2 TPM across the 83 transcriptomes. The raw TPM values were normalized using quantile normalization and rank-based inverse normal transformation [47]. We inferred hidden confounding variables from the normalized gene expression levels using Probabilistic Estimation of Expression Residuals (PEER) [48].

### eQTL mapping

The two VNTR haplotypes (hap1 and hap2) segregating in the Brown Swiss and Original Braunvieh breeds were derived from binned long read alignments with minimap2 (see above). The resulting diplotypes were reconstructed for all bulls of the eQTL cohort based on the Beagle-phased genotypes (see above) for two SNPs (Chr12:86160984 and Chr12:86160971) and one indel (Chr12:86161000) that tagged the VNTR haplotypes. A linear model was fitted using the lm()-function in R to associate normalized gene expression with the VNTR haplotype (coded as 0, 1, and 2 for hap1/hap1, hap1/hap2, and hap2/hap2, respectively) while considering five PEER factors and the top three principal components of the genomic relationship matrix as covariates to account for technical bias and population stratification. An eQTL analysis within the *cis*-regulatory range of genes of interest was conducted using the linear model described above. The VNTR haplotype was fitted as an additional covariate in the conditional analyses. Bonferroni-correction was applied to determine significance thresholds.

### Allelic imbalance analysis

Polymorphic sites overlapping *LOC112449094* exons were extracted from the cohort-level genomic VCF file. Reference and alternate allele support at heterozygous genotypes was subsequently extracted from the WASP-filtered RNA sequencing read alignments using bcftools mpileup. An exact binomial test as implemented in the R binom.test()-function was applied to test for allelic imbalance.

## Declarations

### Ethics approval and consent to participate

Not applicable

### Consent for publication

Not applicable

### Availability of data and materials

Scripts and pipelines used in this work are available online (https://github.com/AnimalGenomicsETH/superpangenome_construction). HiFi reads of two BSW and two OBV cattle are available in the ENA database at the study accession PRJEB42335 under sample accession SAMEA111341789, SAMEA111341790, SAMEA111341791, and SAMEA111341792. DNA and RNA sequencing data of the eQTL cohort are available in the ENA database at the study accession PRJEB46995.

## Authors’ contributions

A.S.L., D.C., and H.P. conceived the study. A.S.L. constructed genome assemblies, constructed/decomposed the pangenomes, and performed the pangenome, variant, and VNTR analyses. D.C. contributed to pangenome construction, pangenome, variant, and VNTR analyses. X.M.M. established the eQTL cohort and aligned the RNA and DNA sequencing data. A.S.L. called variants for the eQTL cohort. H.P. performed the eQTL analyses. M.B. contributed to the design of the eQTL analyses. A.S.L., and H.P. wrote the manuscript with input from D.C. All authors read and approved the final manuscript.

## Acknowledgements

We thank Dr. Anna Bratus-Neuenschwander and Dr. Catharine Aquino from the ETH Zurich technology platform FGCZ (https://fgcz.ch) for DNA fragment analysis and DNA and RNA sequencing.

