## Supplementary_table_figures for "Graph construction method impacts variation representation and analyses in a bovine super-pangenome": Supplementary_files.pdf

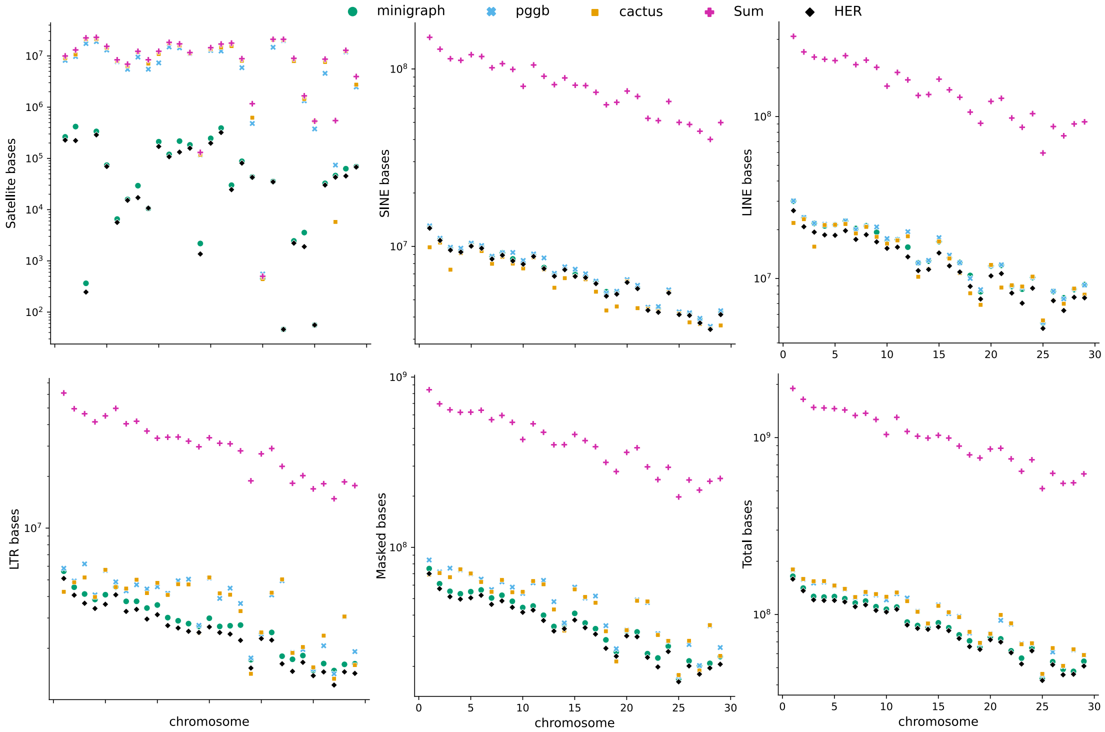


***Supplementary Figure 1****. Repetitive sequence content of the three pangenomes, the sum across all 12 input assemblies (Sum), and the reference genome (HER). Many repetitive elements appear to be compactly represented by the pangenomes, and hence are substantially below the sum of the assemblies. However, centromeric sequences (Satellite) are either too biologically diverged or current mapping/smoothing algorithms are insufficient to collapse sequence from multiple assemblies into fewer nodes. As expected, minigraph contains effectively only centromeric sequence from the reference backbone and is unable to incorporate centromeric sequence that extends beyond the backbone.*

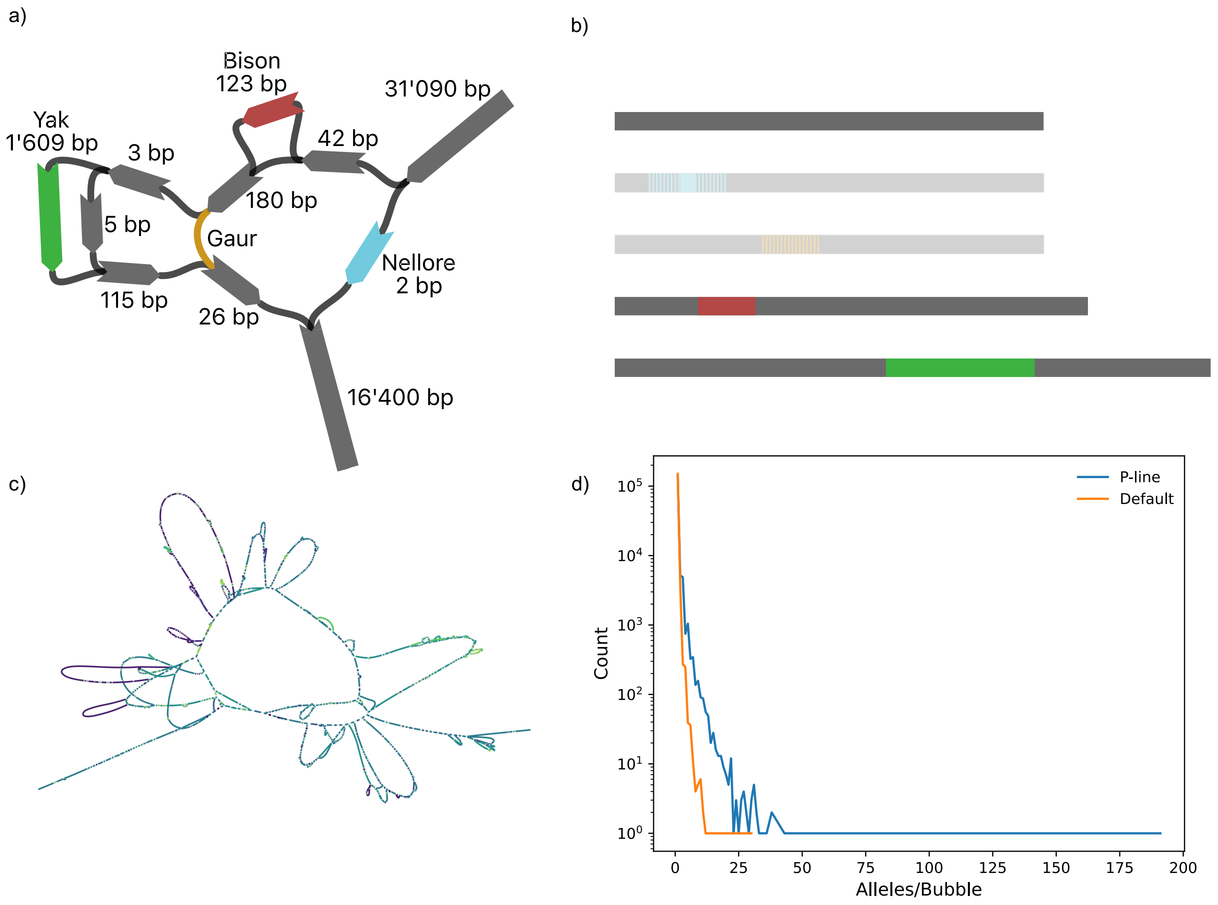


***Supplementary Figure 2.*** *a) Complex bubble in minigraph (BTA29: 2250204-2250575) where reference nodes are dark gray and nonreference nodes (and nonreference deletions) are colored. b) Of the 5 alleles (including reference) observed in the graph, the faded Nellore and Gaur deletions are not recovered when deconstructing the minigraph pangenome without path information into VCF, leaving only 2 insertion and the reference alleles. When path information is added, all 5 alleles are correctly recovered. Minigraph by default does not include P-lines, which has a large detrimental effect on calling SVs in multi-allelic bubbles. c) A complex region (BTA27:6357301-7209140) produces 201 SV alleles without path information, and 462 SV alleles with P-lines. d) Including P-lines greatly increases the number of alleles found per bubble compared to default minigraph with no path information. The effect is magnified for complex bubbles containing many nodes and edges.*

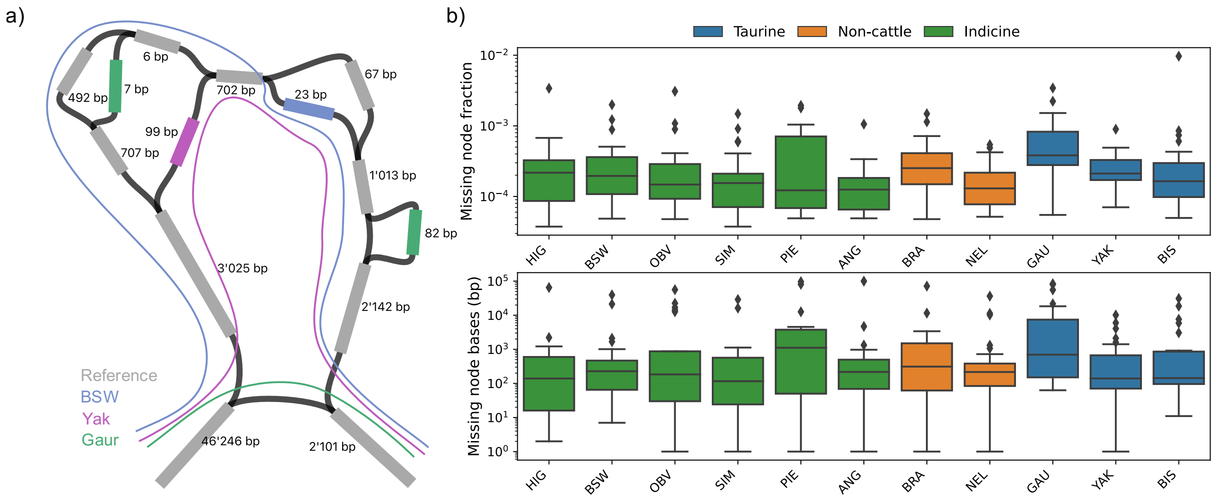


***Supplementary Figure 3.*** *Adding P-lines post-hoc can lead to unexpected realignments. a) An extreme example bubble containing non-reference nodes for BSW, yak, and gaur. Realignment correctly traces BSW and yak through the appropriate non-reference nodes, while gaur is aligned through a deletion, incorrectly tracing the path which should include the two gaur non-reference nodes. As realignment produces an incorrect path, variant calling incorrectly assigns gaur a single 8,091 bp deletion as opposed to 485 bp deletion (top left) and 82 bp insertion (right). b) Some taurine, indicine, and non-cattle nodes included in minigraph pangenomes do not appear in realignment (top), although they do not account for a substantial portion of non-reference sequence found in the graphs (bottom).*

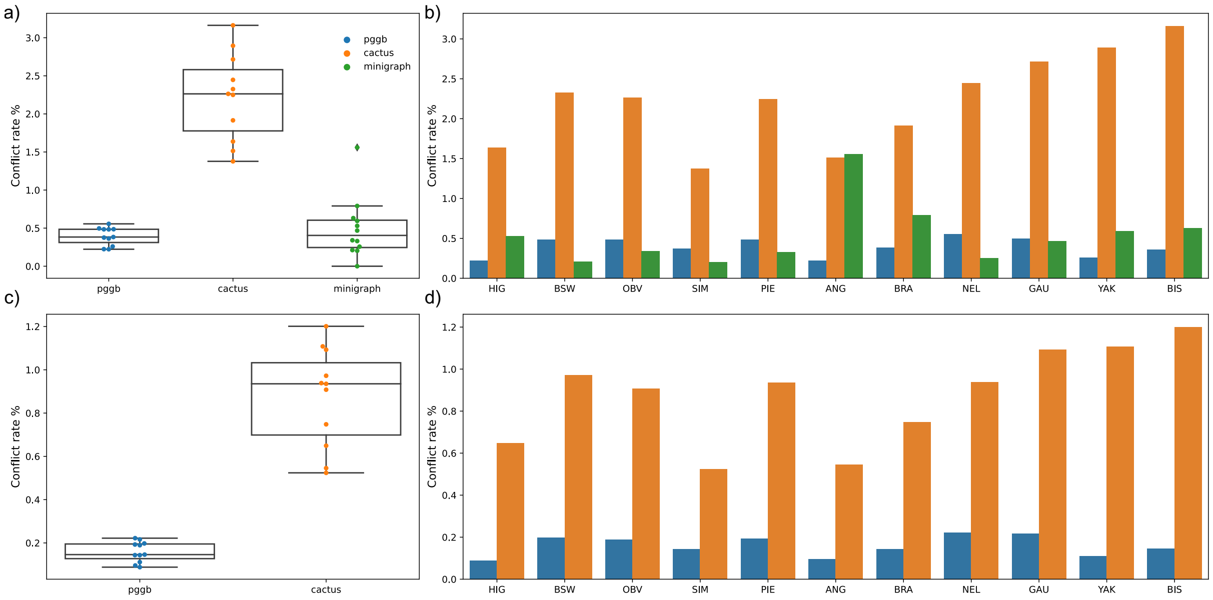


***Supplementary Figure 4****. Proportion of conflicting genotypes detected from the three pangenomes from realigned assemblies for SVs (a) and small variations (c). Each point in the boxplots and every bar indicates the proportion of conflicting genotypes for each assembly. Conflict rates for cactus and pggb were determined by the CONFLICT tag assigned by vg deconstruct. Conflict rates for minigraph were determined during the post hoc realignment to add P-lines, when an assembly had a no call over a bubble. Per-sample conflict rates are shown for SVs (b) and small variation (d).*

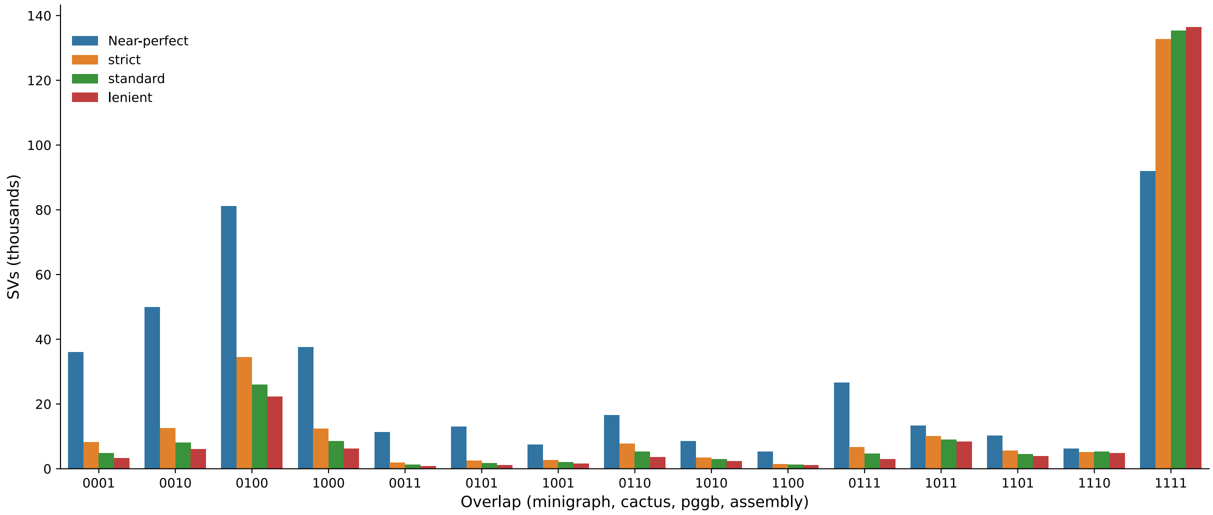


***Supplementary Figure 5****. Overlap of SVs between the three pangenomes and assemblies. Near-perfect overlaps require exact basepair-level resolution and >85% sequence identity to merge SVs, while strict merges variants within 5 bp and >50% sequence identity. Almost all SVs that can be merged at the most lenient level are merged at this strict level, showing most SVs are highly similar between the pangenomes and assemblies.*

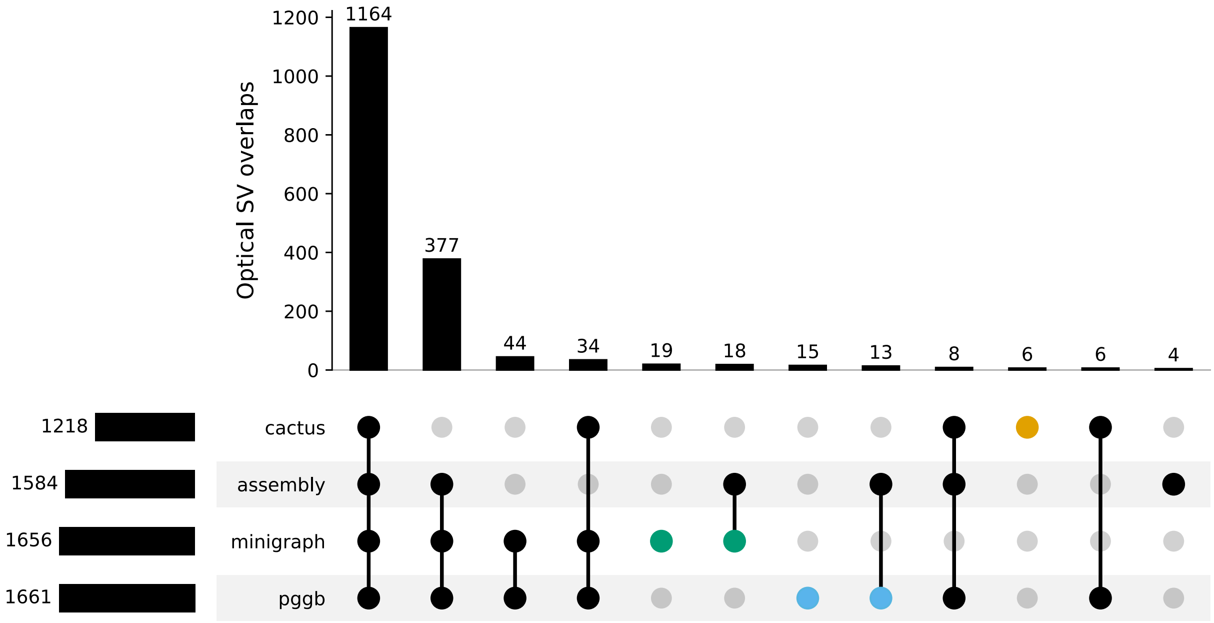


***Supplementary Figure 6.*** *Intersections of the three pangenomes and assembly-derived calls with external optical mapping data. Only variants overlapping with the optical maps are shown. SVs private to one of the three pangenomes (including private to pangenome and assembly) are shown in blue, orange, and green respectively for pggb, cactus, and minigraph.*

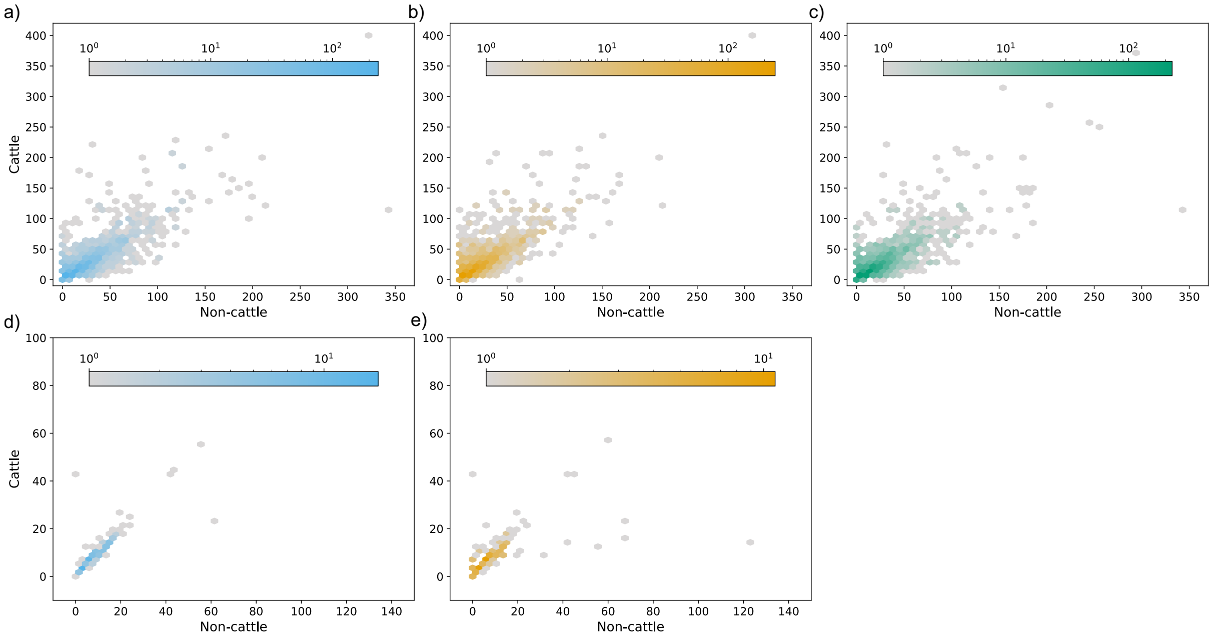


***Supplementary Figure 7**. a), b), c) TR counts that did not agree across all the pangenomes are still broadly similar for pggb, cactus, and minigraph respectively. d), e) Similarly, TRs present in pggb and cactus but not minigraph that disagreed on count are still broadly similar and exhibit low variability between cattle and non-cattle.*

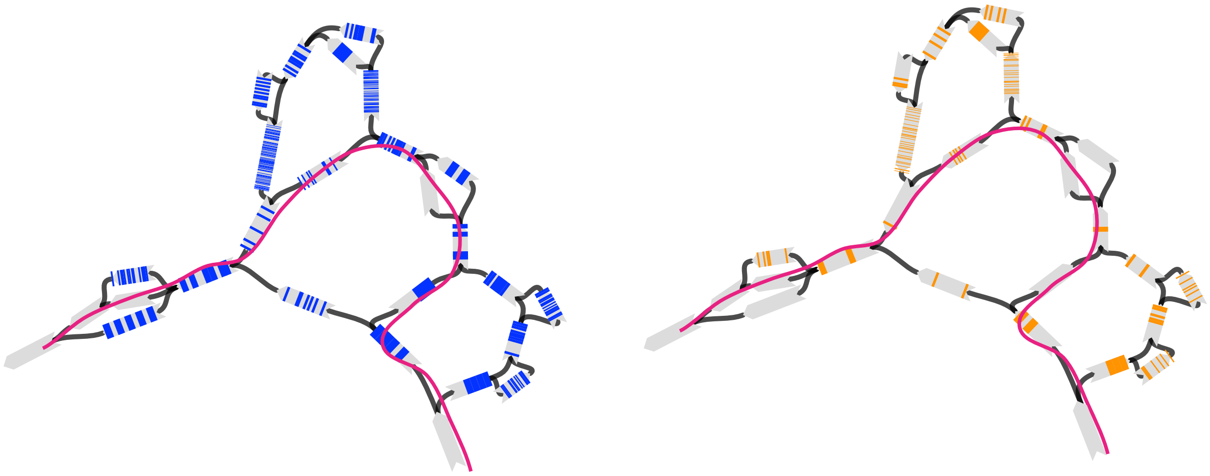

***Supplementary Figure 8.*** *VNTR in minigraph from Figure 6 visualized for two TR motifs separately (blue: TGGATGGGTGGA and orange: TGGGTGGACGGA). The reference path is highlighted in pink. The reference path (generally taken by taurine) contains more of the blue motif, while the non-reference paths contain most of the rarer orange motif. In addition to revealing TR count variation related to evolutionary histories, motif variation is also accessible (albeit to a lesser degree in minigraph) through pangenomes and Bandage plots.*

a

b

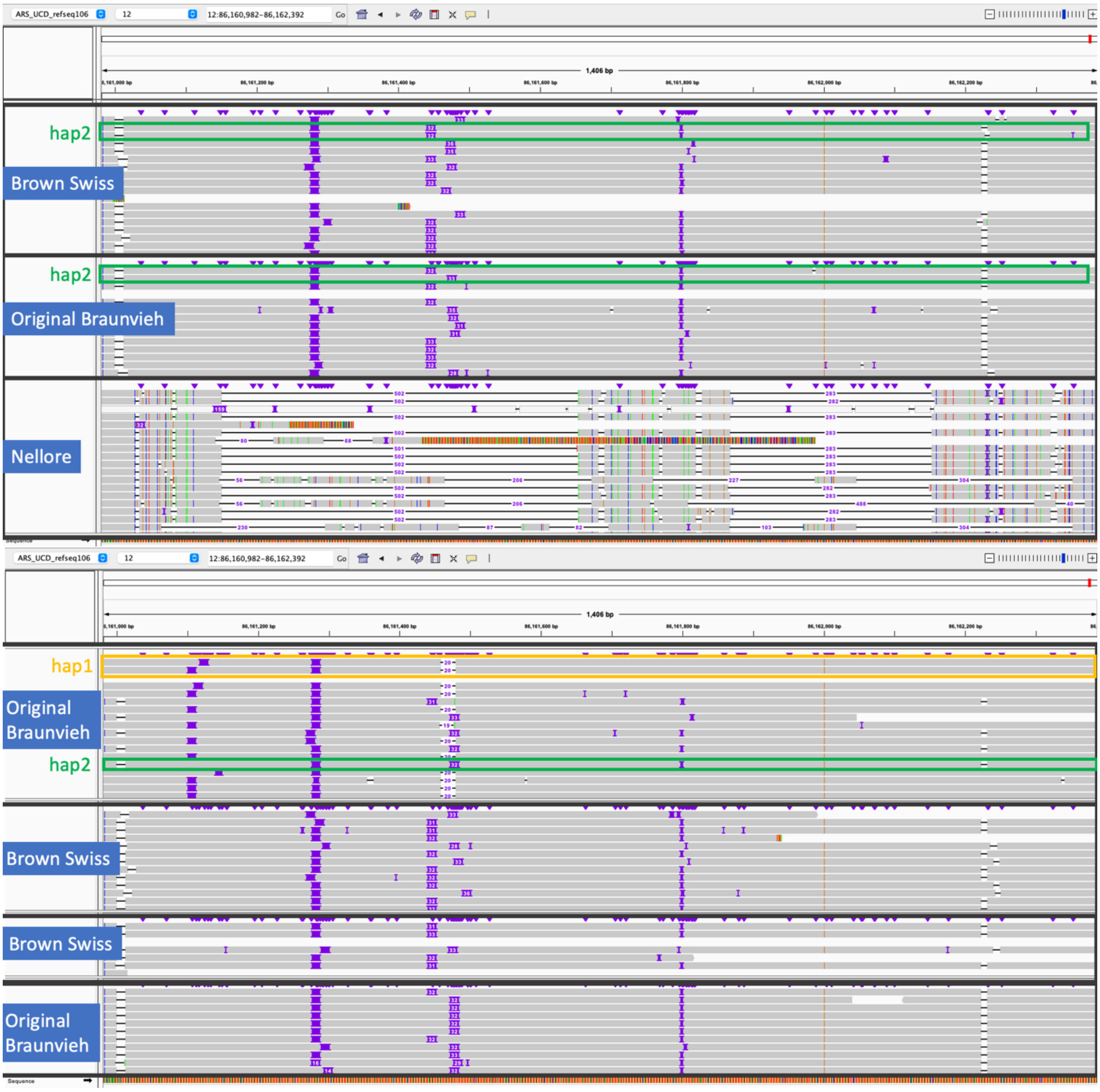


***Supplementary Figure 9**: Detection of two VNTR haplotypes in Original Braunvieh and Brown Swiss cattle. a) IGV screenshots of alignments from parental-binned HiFi reads of Brown Swiss, Original Braunvieh, and Nellore haplotypes that are included in the pangenomes. The screenshots cover the VNTR region (chr12:86,160,982-86,162,392). The Brown Swiss and Original Braunvieh haplotypes that are part of the pangenomes carry haplotype "hap2". The Nellore haplotype is considerably shorted and diverged from the taurine haplotypes. b) We observe a second VNTR haplotype ("hap1") in HiFi read alignments of four additional Original Braunvieh and Brown Swiss animals.*

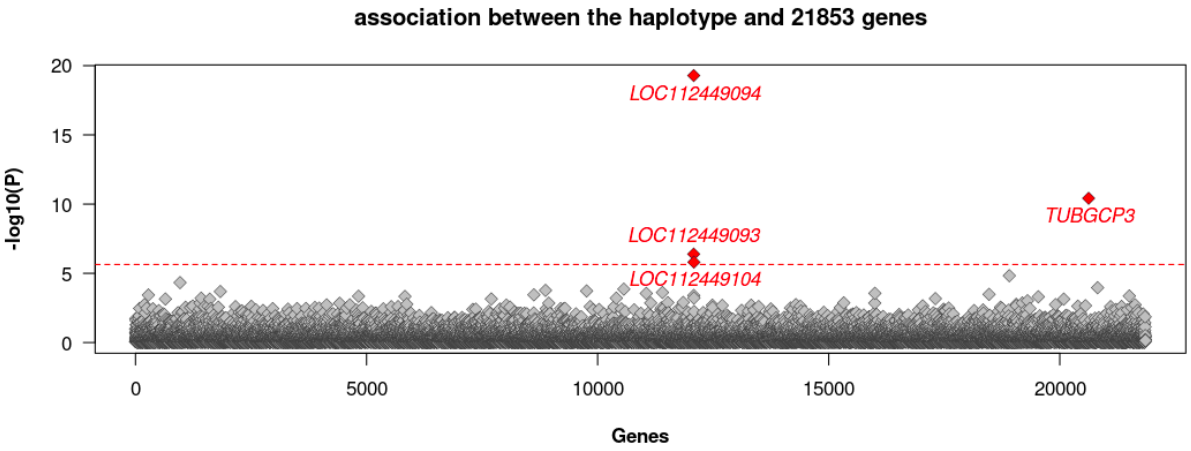


***Supplementary Figure 10***: *eQTL mapping between the VNTR haplotype and normalized expression of 21,853 genes. Genes and ncRNA are ordered in alphabetical order. The red line indicates the Bonferroni-corrected significance threshold (P=0.05 / 21853). Genes and ncRNA exceeding the Bonferroni-corrected threshold are coloured on red.*

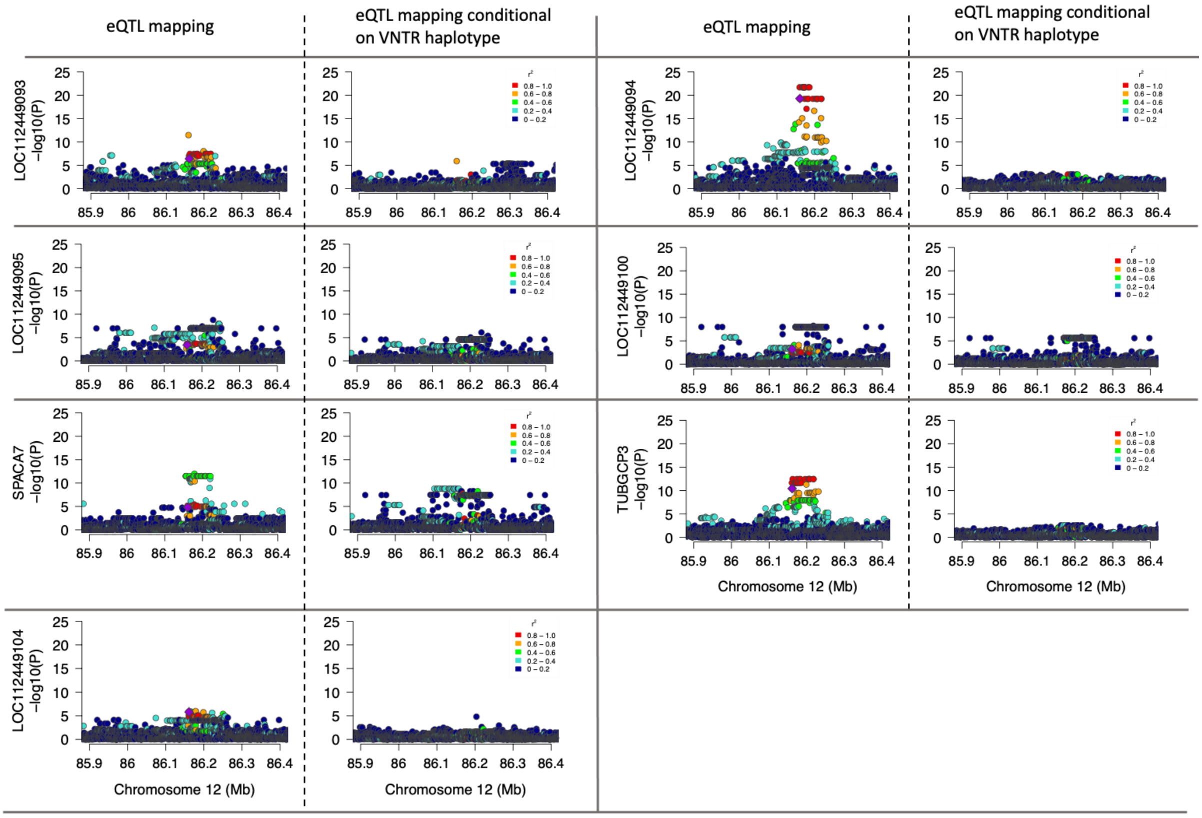


***Supplementary Figure 11***: *cis-expression QTL mapping for two genes and four lncRNA that were significantly associated with the VNTR haplotype. Variants within 1Mb of the genes were tested for association. Different colours indicate the pairwise linkage disequilibrium ($r^2$) between the VNTR haplotype (violet) and all other variants. The cis-eQTL mapping was repeated conditional on the VNTR haplotype.*

***Supplementary Table** 1. Cumulative CPU hours and maximum memory needed for vg deconstruct and the number of variants recovered for the three pangenomes.*

| Tool | CPU hours | Memory (GB) | Variants |
|---|---|---|---|
| minigraph | 0.2 | 0.5 | 164,723 |
| pggb | 10.1 | 11.1 | 57,350,905 |
| cactus | 28.3 | 12.8 | 61,028,922 |

***Supplementary Table** 2. CSV containing the number of centromeric sequence bases trimmed from the start of the chromosomes, bulk sequence bases, and telomeric sequence trimmed from the end for each chromosome and each assembly.*

| |
|---|
| (External file) |

***Supplementary Table 3.** GraphAligner alignments using relaxed (-x dbg -C 100000 --max-trace-count 5 --seeds-minimizer-ignore-frequent 0.001 --precise-clipping 0.9) or strict (-x vg) alignment parameters. The edit rate and query coverage are taken from a single 500 Kb window in chromosome 12 of the Brown Swiss assembly (start at 60867381), chosen for its complexity and subsequent poor relaxed alignment. CPU time is given in seconds and Memory is peak RAM usage in GB. Strict alignment quickly requires more resources for pggb and becomes prohibitive for cactus for whole chromosome alignment.*

| | Relaxed | | | | Strict | | | |
|---|---|---|---|---|---|---|---|---|
| | Edits | Query coverage | CPU | Memory | Edits | Query coverage | CPU | Memory |
| minigraph | 2285 | 99.6 | 50 | 0.7 | 2285 | 99.6 | 99 | 1.0 |
| cactus | 0 | 0 | 164 | 6.9 | 18 | 100 | 244 | 14.1 |
| pggb | 0 | 0 | 140 | 5.1 | 17 | 100 | 189 | 8.1 |

***Supplementary Table** 4. Compute resources for aligning the 12 assemblies included in pangenome construction and the 8 held out for analysis. CPU hours are averaged per assembly for both the bulk and centro-/telomeric regions, and memory is the peak RAM usage across all assemblies.*

| | CPU hours | | | | Memory (GB) | | | |
|---|---|---|---|---|---|---|---|---|
| | Included | | Held-out | | Included | | Held-out | |
| | Bulk | Centro-/telo | Bulk | Centro-/telo | Bulk | Centro-/telo | Bulk | Centro-/telo |
| pggb | 3.7 | 2.4 | 121.1 | 1.8 | 31.7 | 109.8 | 108.2 | 79.6 |
| cactus | 5.6 | 0.9 | 73.2 | 1.3 | 130.4 | 99.4 | 349.5 | 126.9 |
| minigraph | 2.6 | 1.3 | 2.8 | 1.3 | 4.5 | 4.3 | 20.8 | 1.9 |

***Supplementary Table** 5. CSV containing the examined tandem repeats in pggb, cactus, and minigraph. Tandem repeats further examined by adVNTR are also indicated.*

| |
|---|
| (External file) |

***Supplementary Table** 6. Several obvious misalignments found in different pangenomes which affects TR count genotyping. Part of the size difference may be true if there is variation in the number of tandem repeats, but the obvious majority of the large mis-translated regions are due to repetitive regions or spurious graph cycles. Δ bp is the size difference for the lifted-over sample TR coordinates from the original reference TR coordiantes.*

| Pangenome | Sample | Chromosome | Reference coordinates | Sample coordinates | Δ bp |
|---|---|---|---|---|---|
| minigraph | YAK | 5 | 118527963-118528562 | 118396757-118426843 | 29.5 Kb |
| pggb | NEL | 13 | 11057154-11057352 | 16184369-23015882 | 6.8 Mb |
| cactus | NEL | 1 | 121617962-121618026 | 58878820-125260767 | 66.4 Mb |
| cactus | SIM | 4 | 3319408-3319495 | 3310542-113157354 | 109.8 Mb |
| cactus | BIS | 8 | 71781915-71781975 | 23187990-108691932 | 85.5 Mb |
| cactus | BIS | 12 | 6381693-6381754 | 657155-36976359 | 36.3 Mb |
| cactus | GAU | 12 | 6381693-6381754 | 439517-36663196 | 36.2 Mb |
| cactus | YAK | 12 | 6381693-6381754 | 107663-36390629 | 36.3 Mb |

***Supplementary Table** 7. Minigraph pangenomes constructed with different "minimum variant length" parameter (L) values. CPU hours required for pangenome construction increased dramatically for L<10 bp. Warnings refer to the number of "impossible insert" warnings issued during pangenome construction, relating to unsuitable graph topology. Bubbles and Nodes respectively refer to the number of top-level bubbles and nodes present across the autosomes. VNTR overlaps is the number of VNTRs (in total 9,568) that overlap with a graph bubble.*

| L (bp) | CPU hours | Warnings | Bubbles | Nodes | Non-reference bases (Mb) | VNTR overlaps |
|---|---|---|---|---|---|---|
| 50 | 13.45 | 131 | 153,865 | 425,245 | 108.85 | 5,742 |
| 30 | 14.80 | 152 | 220,761 | 611,795 | 110.66 | 6,552 |
| 10 | 15.32 | 222 | 709,032 | 1,977,837 | 118.07 | 7,613 |
| 5 | 29.35 | 484 | 1,571,642 | 4,613,837 | 128.20 | 7,898 |
| 2 | 53.37 | 24,678 | 8,125,250 | 26,623,047 | 220.19 | 8,441 |
